# Proteomics and phosphoproteomics profiling in glutamatergic neurons and microglia in an iPSC model of Jansen de Vries Syndrome

**DOI:** 10.1101/2023.07.08.548192

**Authors:** Jennifer T. Aguilan, Erika Pedrosa, Hedwig Dolstra, Refia Nur Baykara, Jesse Barnes, Jinghang Zhang, Simone Sidoli, Herbert M. Lachman

**Affiliations:** Department of Pathology Albert Einstein College of Medicine 1300 Morris Park Ave. Bronx, NY, 10461; Department of Psychiatry and Behavioral Sciences Albert Einstein College of Medicine 1300 Morris Park Ave. Bronx, NY, 10461; Department of Microbiology & Immunology Albert Einstein College of Medicine 1300 Morris Park Ave. Bronx, NY, 10461; Department of Biochemistry Albert Einstein College of Medicine 1300 Morris Park Ave. Bronx, NY, 10461; Department of Medicine Albert Einstein College of Medicine 1300 Morris Park Ave. Bronx, NY, 10461; Dominick P. Purpura Department of Neuroscience Albert Einstein College of Medicine 1300 Morris Park Ave. Bronx, NY, 10461; Department of Genetics Albert Einstein College of Medicine 1300 Morris Park Ave. Bronx, NY, 10461

**Keywords:** pediatric acute-onset neuropsychiatric syndrome (PANS), autism, regression, POGZ, ubiquitin ligase, UBR4, SRRM1, NUCKS1, PPM1D, Jansen de Vries Syndrome

## Abstract

**Background:** Jansen de Vries Syndrome (JdVS) is a rare neurodevelopmental disorder (NDD) caused by gain-of-function (GOF) truncating mutations in *PPM1D* exons 5 or 6. PPM1D is a serine/threonine phosphatase that plays an important role in the DNA damage response (DDR) by negatively regulating TP53 (P53). JdVS-associated mutations lead to the formation of a truncated PPM1D protein that retains catalytic activity and has a GOF effect because of reduced degradation. Somatic *PPM1D* exons 5 and 6 truncating mutations are well-established factors in a number of cancers, due to excessive dephosphorylation and reduced function of P53 and other substrates involved in DDR. Children with JdVS have a variety of neurodevelopmental, psychiatric, and physical problems. In addition, a small fraction has acute neuropsychiatric decompensation apparently triggered by infection or severe non-infectious environmental stress factors.

**Methods:** To understand the molecular basis of JdVS, we developed an induced pluripotent stem cell (iPSC) model system. iPSCs heterozygous for the truncating variant (*PPM1D*^+/tr^), were made from a patient, and control lines engineered using CRISPR-Cas9 gene editing. Proteomics and phosphoprotemics analyses were carried out on iPSC-derived glutamatergic neurons and microglia from three control and three *PPM1D*^+/tr^ iPSC lines. We also analyzed the effect of the TLR4 agonist, lipopolysaccharide, to understand how activation of the innate immune system in microglia could account for acute behavioral decompensation.

**Results:** One of the major findings was the downregulation of POGZ in unstimulated microglia. Since loss-of-function variants in the *POGZ* gene are well-known causes of autism spectrum disorder, the decrease in *PPM1D*^+/tr^ microglia suggests this plays a role in the neurodevelopmental aspects of JdVS. In addition, neurons, baseline, and LPS-stimulated microglia show marked alterations in the expression of several E3 ubiquitin ligases, most notably UBR4, and regulators of innate immunity, chromatin structure, ErbB signaling, and splicing. In addition, pathway analysis points to overlap with neurodegenerative disorders.

**Limitations:** Owing to the cost and labor-intensive nature of iPSC research, the sample size was small.

**Conclusions:** Our findings provide insight into the molecular basis of JdVS and can be extrapolated to understand neuropsychiatric decompensation that occurs in subgroups of patients with ASD and other NDDs.

## Introduction

Jansen de Vries Syndrome (JdVS) (OMIM 617450) is a recently discovered neurodevelopmental disorder (NDD) caused by truncating mutations in *PPM1D* exons 5 or 6 (1–4). It is characterized by mild to severe intellectual disability, anxiety disorder, attention deficit hyperactivity disorder (ADHD), obsessive behavior, hypotonia, sensory integration problems, and in some cases, autism spectrum disorder (ASD). In addition, feeding difficulties and gastrointestinal problems (e.g., constipation, esophageal reflux, and cyclic vomiting syndrome) are common. Approximately half of the reported cases have an increase in childhood infections, although this has not been systematically evaluated and the pathogenesis has not been established. *PPM1D* codes for a member of the PP2C serine/threonine phosphatase family. So far, every *PPM1D* mutation found in JdVS is predicted to translate into a truncated protein (e.g., nonsense mutations and frameshifts) because of the loss of C-terminal amino acids. The catalytic domain encoded largely by exons 1-4 is preserved, and an increase in PPM1D half-life occurs because truncated proteins lose a degradation signal that maps within the terminal 65 amino acids (5,6).

*PPM1D* is a well-known tumor suppressor gene, acting as a negative regulator of P53 and other proteins involved in the DNA damage response (DDR) pathway, such as MDM2, ATM, CHK1, CHK2, ATR, and H2AX (7–9). Somatic GOF truncating mutations in exons 5 or 6 have been found in a variety of cancers (7–14). Cancer risk in JdVS has not yet been established, although a normal P53 response to ionizing radiation was found in EB-transformed lymphocytes derived from children with JdVS (4).

In addition to the neurodevelopmental and psychiatric features of JdVS, a small subgroup of patients experience behavioral decompensation that appears to be linked to infection or severe physical stress. One patient we identified was diagnosed with pediatric acute-onset neuropsychiatric syndrome (PANS) as a child, several years prior to exome sequencing for NDD revealed a typical *PPM1D* truncating variant (15). PANS is an enigmatic, neuroinflammatory disorder characterized by the abrupt onset of severe neurological and psychiatric symptoms that includes obsessive-compulsive disorder (OCD), restricted eating, anxiety, cognitive deficits with academic regression, disrupted sleep, rage, mood disturbance, joint inflammation, and autonomic nervous system disturbances (e.g., enuresis, postural orthostatic tachycardia syndrome) (16–18). Subsequently, we identified two other JdVS case in which severe behavioral and motor regression occurred following infection and noninfectious triggers.

Mouse *Ppm1d* knockout (KO) models have been developed, which show effects on dendritic spine morphology and memory processes, a disturbance in T- and B-lymphocyte differentiation, proliferation, cytokine production, and an increase in phagocytosis and autophagy in peripheral macrophages (19–22). However, a mouse *Ppm1d* KO is not an appropriate model for JdVS GOF variants. Consequently, in order to understand the underlying molecular basis of truncated PPM1D on neuronal function and the apparent propensity a subgroup of JdVS patients has for acute neuropsychiatric decompensation, we developed an induced pluripotent stem cell (iPSC) model and analyzed glutamatergic neurons and microglia by proteomics and phosphoproteomics.

## Methods

### Subjects

The JdVS patient is a male who was born full-term following an uncomplicated pregnancy who was diagnosed as a child following whole exome sequencing (WES), which revealed a typical *PPM1D* truncating mutation in exon 5 (c.1210C>T; p.Q404X). All *PPM1D* heterozygotes, whether patient-derived or developed using CRISPR-Cas9 editing (see below), will be referred to as *PPM1D*^+/tr^. Analysis of parental DNA showed that the mutation was de novo, as is the case for >90% of JdVS cases. His typically developing brother was used as one of the controls. Another typically developing control male was used to develop isogenic iPSC *PPM1D*^+/tr^ lines in which a truncating mutation was introduced in exon 5 using CRISPR-Cas9 gene editing (see **Additional file 1: Expanded Methods** for details). A third male was used as a typically developing control. These last two controls were characterized in another study (23).

### Development of iPSCs from peripheral blood CD34+ cells

All methods used in this study are described briefly here; details can be found in **Additional file 1: Expanded Methods**. iPSC lines were generated from human peripheral blood CD34+ hematopoietic stem cells (HSC) with a CytoTune-iPS 2.0 Sendai Reprogramming Kit (Invitrogen) following the manufacturer’s protocol, as previously described (24). All lines were capable of differentiating into the three germ layers and showed no expression of reprogramming transcription factors. Cytogenetic analysis was negative.

### CRISPR-Cas9 gene editing

A heterozygous truncating variant in *PPM1D* exon 5 was generated by CRISPR-Cas9 gene editing, using a protocol described by Ran et al (25). Briefly, a guide RNA (gRNA) sequence coding for a region in exon 5 adjacent to a PAM sequence 9 base pairs from the patient mutation was chosen (**Figure 1**: **Additional file 1: Expanded Methods**).

**Figure 1.**
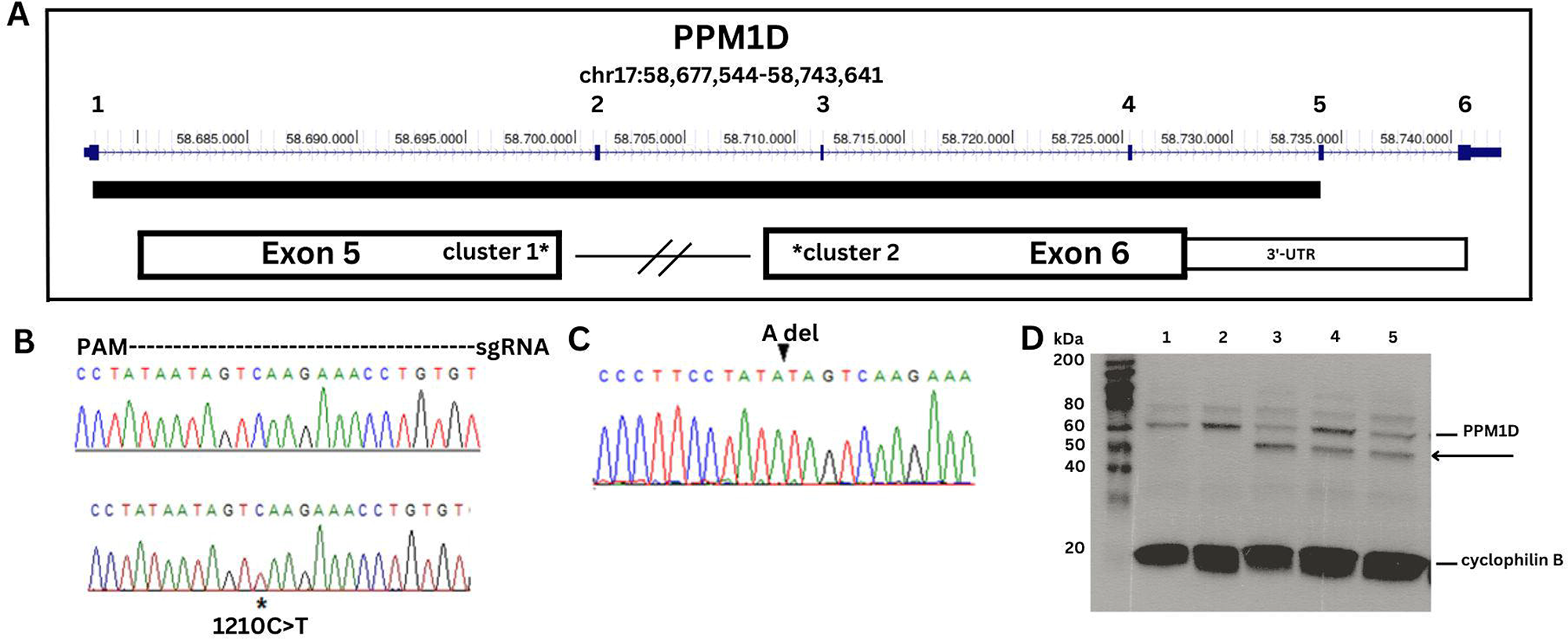
DNA sequence analysis and Western blot of PPM1D truncating variants. A. Map of PPM1D showing the 6 exons with the catalytic domain depicted as a solid bar. The two clusters of JdVS mutations are in the 3’-end of exon 5 and the 5’-end of exon 6. B. DNA sequence strip of wild type allele on top and the patient sample showing the c.1210C>T; p.Q404X nonsense mutation on bottom. The region covered by the guide RNA used for CRISPR-Cas 9 engineering is shown. C. Two isogenic lines with an “A” deletion were generated with CRISPR. D. PPM1D Western blot showing wild-type protein and the truncated protein. Cyclophilin is a loading control. Lanes 1 and 2 are control samples, lane 3 is the patient sample, and lanes 4 and 5 are two CRISPR lines.

### Differentiation of iPSCs into glutamatergic neurons

iPSCs were maintained as previously described (23). Glutamatergic neuronal differentiation was induced using a protocol developed by Zhang et al., in which differentiation is driven by overexpression of the transcription factor NGN2 (26). A tet-inducible expression system was introduced and lentivirus particles prepared from the plasmid vectors; pLV_TRET_hNgn2_UBC_Puro (plasmid #61474) and FUdeltaGW-rtTA (plasmid #19780), followed by treatment with doxycycline and selection with puromycin (see **Additional file 1: Expanded Methods** for details). The protocol routinely leads to the production of a nearly pure culture of excitatory cortical glutamatergic neurons.

### Neurite outgrowth

Neurite outgrowth was assessed blind to genotype using the NeuronJ plugin (27). Glutamatergic neurons were stained with Map2 and Tuj1 antibodies and imaged at 10x resolution. Images of patient and control neurons were converted to 8-bit grayscale, and individual dendrites were traced and labeled using the semiautomatic manual tracing tool. Approximately 10 images with an average of 12 neurons per field were analyzed per sample. The “measure tracings” function was used to determine mean length (in pixels) of the dendritic branches.

### Differentiation of iPSCs into microglia

To generate microglia, we used kits from STEMCELL^TM^ Technologies (STEMdiff^TM^ Hematopoietic Kit, catalog number 05310; STEMdiff^TM^ Microglia Differentiation Kit, catalog number 100-0019; STEMdiff^TM^ Microglia Maturation Kit, catalog number 100-0020) according to the manufacturer’s instructions, with minor modifications as described in **Additional file 1: Expanded Methods**. iPSCs are first differentiated into HSCs, followed by terminal differentiation into microglia. The microglia grow in suspension with the control and PPM1D^+/tr^ showing a similar morphology. (**Additional file 2: Fig. S1**).

### Fluorescence-activated cell sorting (FACS)

Single-cell suspensions were used for flow cytometry staining. We followed a protocol for Staining Cell Surface Targets for Flow Cytometry from ThermoFisher. All antibodies were obtained from Stemcell Technologies, except for TMEM119, which is from Novus Biologicals. For HSCs, we used CD45 FITC (Catalog number 60018FI.1) CD43 APC (Catalog number 60085AZ.1), and CD34 PE (Cat. 60013PE.1) antibodies. For microglia we used TMEM119 APC (Catalog FAB10313A) and CD11b PE (Catalog 60040PE.1) antibodies. Antibody concentrations were 5ul per 100ul for all Ab except TMEM119 for which 0.5ul per 100ul was used. Flow cytometry acquisition was obtained using a BD LSRII analyzer, and BD FlowJo software was used for data analysis.

### Cytokine array

Microglia were seeded at 5 x 10^5^ cells/well in a 12 well, Matrigel-coated plate in STEMdiff Microglia Maturation media 24 days post differentiation. After 5 days of maturation, cells were stimulated with 100ng/ml LPS (O111:B4 strain; Sigma catalog # L4391) for 24 hours at 37°C. Supernatants were collected and analyzed using the Proteome Profiler Array Human Cytokine Array (R&D Systems catalog # ARY005B) following the manufacturer’s instructions. Arrays were analyzed using Quick Spots Image Analysis Software. Each cytokine and chemokine on the array is measured in duplicate.

### Western Blotting

Proteins were prepared with Pierce^TM^ RIPA Buffer (Thermoscientific catalog # 89900) according to the manufacturer’s protocol, with a protease inhibitor cocktail mix (Sigma catalog # P8340). Protein concentrations were verified using the BCA assay. Western Blotting was essentially carried out as previously described, with modifications, as described in **Additional file 1: Expanded Methods**. Phosphorylation of CaMKII (CaMK2) was analyzed by comparing the phosphorylated and unphosphorylated proteins, which are also described in more detail in the expanded methods section.

### Proteomics and phosphoproteomics

Proteomics and phosphoproteomics, and subsequent bioinformatics analyses were performed as previously described (28–35) (see **Additional file 1: Expanded Methods** for details)

## Results

### Development of iPSCs

A patient-specific line was developed from a male with JdVS who had a de novo nonsense mutation at codon 404 in exon 5 (c.1210C>T; p.Q404X) (**Figure 1**). The same mutation was found in one of the subjects in the original JdVS paper (individual 4) {{6152 Jansen,S. 2017}}. His typically developing brother was used as a control. We also used CRISPR-Cas9 gene editing on another control line to create truncating mutations in exon 5 near the patient’s variant. Two clones with an “A” deletion 5 bp from the patient mutation were obtained. The deletion causes a frameshift and premature termination after 6 additional amino acids are inserted (c.1209delA; N402Ifs*6). This is still within the boundaries of the most proximal truncating mutation described by Jansen et al., at cDNA position 1188 (4). Both the patient sample and the CRISPR-engineered lines show the truncated protein on a Western blot (**Figure 1**).

### Proteomics: glutamatergic neurons

Proteomics and phosphoproteomics were carried out on glutamatergic neurons (day 21] differentiated from iPSCs. A total of four control and five PPM1D+/tr samples were analyzed. 4,948 proteins were detected among which 35 were significantly upregulated and 26 that were downregulated in the *PPM1D*^+/tr^ neurons (p<0.05, corresponds to a p-value of 4.32, see **Additional file 3: Table S1**). The volcano plots for this analysis, as well as the subsequent proteomics and phosphoproteomics data described below are shown in **Figure 2**.

**Figure 2.**
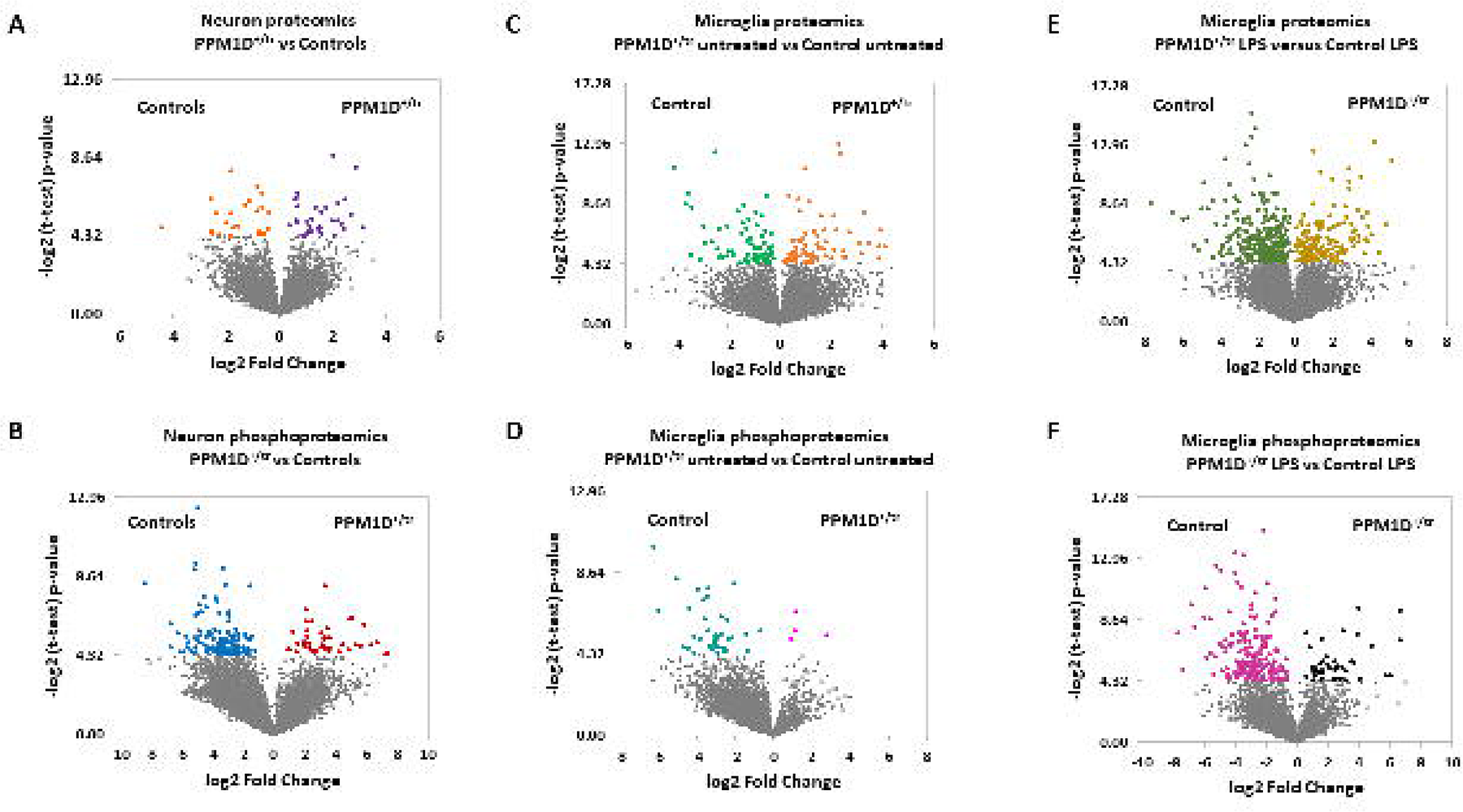
Volcano plots of differentially expressed proteins and phosphoproteins for all analyses. -log2 value of 4.32 corresponds to p=0.05, which is the cutoff.

Gene Ontology (GO) analysis was carried out to characterize the pathways and processes affected by differentially expressed proteins (DEPs). The top GO pathway was, surprisingly, positive regulation of T cell differentiation (**Table 1; Additional file 3: Table S1**). This is probably due to expression of regulatory factors influenced by PPM1D that are expressed in both neurons and T-cells, an idea supported by the finding that DEPs contributing to the T cell differentiation GO term in neurons; CBFB, PNP, AP3D1, ANXA1, AP3B1, SART1, BAD, STAT5B, and ZMIZ1, are also expressed in peripheral blood mononuclear cell (PBMC) types and microglia (36). In addition, Ppm1d has been found to regulate T_h_9 cell development and T-cell differentiation in mice (37,38). Thus, the common regulation of these proteins in neurons and immune cells could be coincidental. The other top GO terms are related to the apparently novel effect of PPM1D on processing H/ACA snoRNAs, a class of small nucleolar RNAs (snoRNAs) that regulate ribosome biogenesis and alternative splicing (39).

**Table 1.** Gene Ontology (GO) analysis, neurons. All differentially expressed proteins and phosphoproteins were used to determine GO terms. Only the most significant terms are shown. For the complete list, see **Additional file 3: Table S1. Tab1.**

| GO term | Neuron Proteomics: Biological Processes | P-value |
| --- | --- | --- |
| GO:0045582 | positive regulation of T cell differentiation | 7.36E-06 |
| GO:0000495 | box H/ACA snoRNA 3'-end processing | 1.30E-05 |
| GO:0034964 | box H/ACA snoRNA processing | 1.30E-05 |
| GO:0000375 | RNA splicing, via transesterification reactions | 2.09E-05 |
| GO:0031126 | snoRNA 3'-end processing | 2.31E-05 |
| GO:0006397 | mRNA processing | 2.77E-05 |
| GO:0000377 | RNA splicing, via transesterification with bulged adenosine as nucleophile | 3.12E-05 |
| GO:0000398 | mRNA splicing, via spliceosome | 3.12E-05 |
| GO:0043144 | snoRNA processing | 3.33E-05 |
| GO:0033979 | box H/ACA snoRNA metabolic process | 3.81E-05 |
| GO:0090669 | telomerase RNA stabilization | 3.81E-05 |
| GO:0045580 | regulation of T cell differentiation | 8.72E-05 |
| GO:0045621 | positive regulation of lymphocyte differentiation | 8.98E-05 |
| GO:0016074 | snoRNA metabolic process | 9.86E-05 |

| GO term | Neuron Phosphoproteomics: Biological Processes | P-value |
| --- | --- | --- |
| GO:0007010 | cytoskeleton organization | 3.52E-07 |
| GO:0071840 | cellular component organization or biogenesis | 2.13E-06 |
| GO:0016043 | cellular component organization | 2.59E-06 |
| GO:0006397 | mRNA processing | 4.87E-06 |
| GO:0016071 | mRNA metabolic process | 7.23E-06 |
| GO:0050684 | regulation of mRNA processing | 6.40E-05 |
| GO:0008380 | RNA splicing | 7.29E-05 |
| GO:0009987 | cellular process | 7.29E-05 |
| GO:1903311 | regulation of mRNA metabolic process | 1.03E-04 |
| GO:1901879 | regulation of protein depolymerization | 1.18E-04 |
| GO:0006396 | RNA processing | 1.60E-04 |

We also analyzed neuronal DEPs by KEGG (Kyoto Encyclopedia of Genes and Genomes), which showed that the top pathway for upregulated proteins was spliceosome, consistent with the GO terms (**Table 2**). In addition, enrichment for proteins involved in several neurodegenerative disorders was also found (e.g., Amyotrophic Lateral Sclerosis [ALS], Huntington’s Disease (HD), Parkinson’s Disease (PD), Alzheimer’s Disease (AD), and prion disease). The top differentially expressed down-regulated KEGG pathways were metabolic pathways, ribosomes, and, similar to the up-regulated pathways, ALS, PD, and HD. These findings suggest that features underlying the pathogenesis of JdVS are shared with those involved in some neurodegenerative disorders.

**Table 2.** KEGG analysis, neurons. KEGG analysis based on differentially expressed up regulated proteins and phosphoprotein, and differentially expressed down-regulated proteins and phosphoproteins. See **Additional file 3: Table S1** for complete lists.

| **KEGG pathway: up-regulated neuronal proteins** | **p-value** |
| --- | --- |
| Spliceosome | 1.56E-25 |
| Amyotrophic lateral sclerosis | 1.48E-14 |
| Nucleocytoplasmic transport | 2.39E-14 |
| Endocytosis | 1.87E-13 |
| Huntington disease | 5.11E-13 |
| Salmonella infection | 9.37E-13 |
| Parkinson disease | 9.78E-13 |
| Alzheimer disease | 6.35E-11 |
| Prion disease | 8.90E-11 |
| Pathways of neurodegeneration - multiple diseases | 9.66E-11 |

| **KEGG pathway: up-regulated neuronal phosphoproteins** | **p-value** |
| --- | --- |
| ErbB signaling pathway | 3.18E-08 |
| Axon guidance | 5.53E-08 |
| Neurotrophin signaling pathway | 2.06E-06 |
| Regulation of actin cytoskeleton | 2.33E-06 |
| Pathogenic Escherichia coli infection | 6.54E-05 |
| Insulin signaling pathway | 8.58E-05 |
| Spliceosome | 4.57E-04 |
| Focal adhesion | 4.57E-04 |
| Autophagy | 5.31E-04 |
| Adherens junction | 6.21E-04 |

| **KEGG pathway: down-regulated neuronal proteins** | **p-value** |
| --- | --- |
| Metabolic pathways | 1.69E-13 |
| Ribosome | 3.29E-13 |
| Amyotrophic lateral sclerosis | 2.01E-09 |
| Parkinson disease | 1.36E-08 |
| Pathways of neurodegeneration - multiple diseases | 4.91E-08 |
| Thermogenesis | 1.33E-07 |
| Huntington disease | 1.96E-07 |
| Oxidative phosphorylation | 4.27E-07 |
| Valine, leucine and isoleucine degradation | 6.49E-07 |
| Endocytosis | 6.74E-07 |

| **KEGG pathway: down-regulated neuronal phosphoproteins** | **p-value** |
| --- | --- |
| Axon guidance | 1.40E-07 |
| Insulin signaling pathway | 1.49E-05 |
| ErbB signaling pathway | 3.74E-05 |
| Regulation of actin cytoskeleton | 3.94E-05 |
| Spliceosome | 1.76E-04 |
| Endocytosis | 6.98E-04 |
| Bacterial invasion of epithelial cells | 1.72E-03 |
| Inositol phosphate metabolism | 2.19E-03 |
| Focal adhesion | 2.19E-03 |
| Phosphatidylinositol signaling system | 1.09E-02 |

Analysis of the top individual up and down-regulated DEPs was particularly noteworthy for altered expression of proteins involved in ubiquitin signaling (**Table 3**). The top-upregulated protein, for example, was CUL4B, a scaffold protein of the CUL4B-Ring E3 ligase complex, which is expressed primarily in the nucleus where it plays a role in DNA repair and tumor progression (40–42). Loss of function (LOF) variants have been found in NDDs (43–46). CUL4B is also an immune regulator, and is involved in the degradation of SIN1, an mTORC2 component (40,47–49).

**Table 3.**
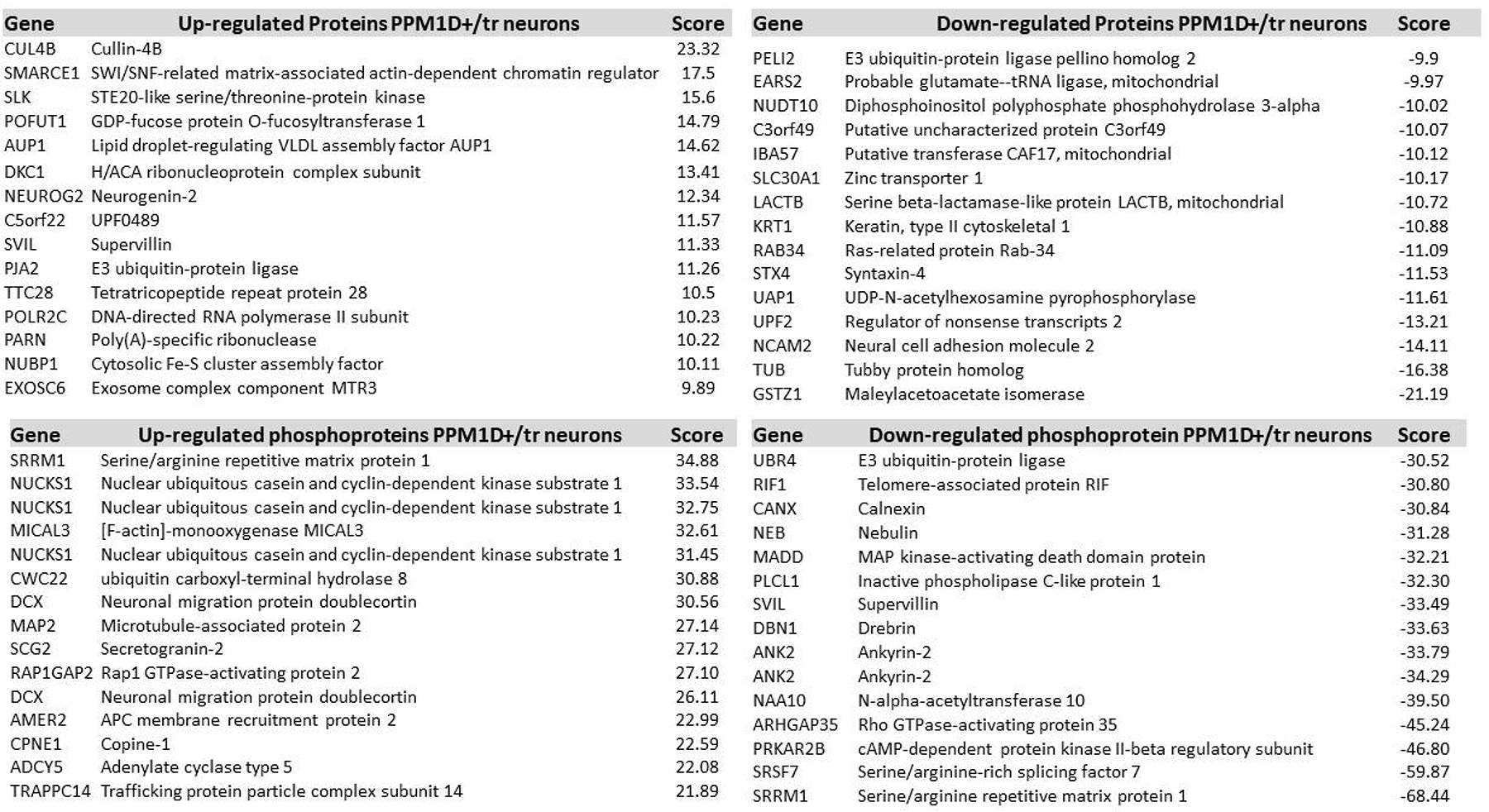
Differentially expressed proteins and phosphoproteins, neurons. Differentially expressed up and down-regulated proteins and phosphoproteins based on total scores (calculated by multiplying Fold Change by the p-value [-log 2 of 4.32 corresponds to p=0.05]). DEPs shown in descending order.

Other ubiquitin signaling proteins among the top 10 upregulated DEPs were PJA2, an E3 ubiquitin-protein ligase, and AUP1, which forms a complex with the ubiquitin-conjugating enzyme (E2), UBE2G2 (50–52). Among the top downregulated DEPs affecting ubiquitin signaling is PELI2, a member of the E3 ubiquitin ligase family that regulate the innate immune system by increasing NLRP3 inflammasome activation (53).

Other top upregulated DEPs of interest include SMARCE1, SLK, POFUT1, DKC1, and NEUROG2. SMARCE1 codes for an SWI/SNF chromatin remodeling complex component that regulates gene expression and can cause ASD when mutated (54–57).

Other proteins that were most downregulated in *PPM1D*^+/tr^ neurons were UPF2, NCAM2, TUB, and GSTZ1. UPF2 is a regulator of nonsense-mediated decay (NMD) and low expression is a factor in resistance to ATR inhibitors: ATR is a DNA damage sensor and a PPM1D substrate (9,58). Disruption of NMD has been associated with neurodevelopmental disorders (59) GSTZ1 is a member of the glutathione S-transferase super-family that detoxifies products of oxidative stress, a process linked to PD, AD, and ALS (60–63).

### Phosphoproteomics: glutamatergic neurons

Since PPM1D is a serine/threonine phosphatase, we also carried out a phosphoproteomics analysis on the samples used in the glutamatergic neuronal proteomics experiment. However, one sample was omitted for technical problems, so 4 control vs 4 *PPM1D*^+/tr^ neuronal samples were analyzed. A total of 7,542 phosphosites were detected (**Additional file 4: Table S2**). At p < 0.05, 174 differentially expressed phosphosites (DEPP) differed significantly between control and PPM1D^+/tr^ neurons; 46 were higher and 128 were lower. GO analysis showed that the most enriched phosphorylations are related to cytoskeleton organization, cellular component organization or biogenesis, and mRNA processing (**Table 1**). KEGG analysis of differentially expressed upregulated proteins showed that the top pathways were ErbB signaling, axon guidance, neurotrophin signaling, and regulation of the actin cytoskeleton (**Table 2**). Axon guidance and ErbB signaling were also among the top downregulated pathways, along with insulin signaling and spliceosome.

The top DEPP that increased in the *PPM1D*^+/tr^ neurons was SRRM1, which is involved in RNA processing, as a component of pre- and post-splicing multiprotein mRNP complexes that play major roles in RNA metabolism (**Table 3**) (63). Altered expression affects prostate cancer aggression and invasion of hepatocellular carcinoma cells (64,65). Mutations in *PPM1D* mutations are associated with both, suggesting that altered SRRM1 phosphorylation plays a role in PPM1D-associated cancers (68, 69). Strikingly, two phosphosites on SRRM1 (Ser725 and Thr727) were also the top downregulated DEPPs. Predicted targets at Thr727 include HIPK1 and p38MAPK (http://www.phosphonet.ca/), which are PPM1D substrates. The findings suggest that regulation of SRRM1 is a novel feature of truncating *PPM1D* variants.

Interestingly, three of the top DEPPs were found in NUCKS1, a chromatin regulator that regulates DNA repair (66–68). NUCKS1 has been implicated in PD in genome wide association studies (GWAS) and is a known PPM1D substrate, although neither of the top three neuronal NUCKS1 phosphosites occurs at SQ/TQ motifs, which are canonical PPM1D targets (9,69,70). Another protein that scored multiple hits among the top upregulated phosphosites is DCX (doublecortin), a cytoskeletal protein that stabilizes microtubules and regulates neuronal migration and cortical layering during development (71).

Differential phosphorylation of proteins that are known PPM1D substrates, such as ATM, CHK1, CHK2, and P53, were not detected, perhaps because the neurons were postmitotic and not subjected to conditions that would most effectively induce their phosphorylation (e.g., ionizing radiation). In fact, among the DEPPs that showed a decrease in phosphorylation expected of a PPM1D GOF effect, only three, ENAH, AKAP12, and ANK2 occurred at SQ sites, suggesting that the majority of neuronal DEPPs are secondary to the downstream effects of PPM1D on other kinases and phosphatases, although novel, noncanonical targets are possible as well.

Overall, the neuronal proteomics and phosphoproteomics data showed differential expression of proteins and phosphoproteins coregulated in T-cells, splicing, DDR, chromatin regulation, neurodegeneration, ErbB signaling, and ubiquitin ligases.

### Proteomics: Microglia

As described in the introduction, we identified several JdVS cases in whom severe motor and behavioral regression occurred following infections and non-infectious stressors. Although these examples of acute neuropsychiatric decompensation appear to be rare occurrences in JdVS, we extended the proteomics analysis to include microglia. An additional rationale is that microglia have been implicated in the pathogenesis of ASD and NDD (72–75). Microglia were developed from six iPSC lines (three control and three *PPM1D*^+/tr^). The differentiation protocol produced similar populations of TMEM119/CD11B double-positive cells; between 71.9 to 88% (**Figure 3**). 5,759 proteins were detected, which included 76 that were significantly upregulated and 76 that were downregulated (**Additional file 5: Table S3; Figure 2**). GO analysis showed enrichment of DEPs related to blood vessel development: aorta morphogenesis, blood vessel lumenization, and blood vessel morphogenesis, although the p-values are modest (**Table 4**). Nevertheless, these findings are of interest. The upregulated proteins that contributed to these GO findings, DLL4, RBPJ, and LRP, are all involved in endothelial function that can affect the brain-blood barrier (BBB) suggesting that *PPM1D* truncating mutations increase BBB permeability (76,77). In fact, PPM1D has been shown to be a BBB regulator (78). The findings support the idea that patients with JdVS are prone to neuroinflammation in response to a peripheral immune challenge.

**Figure 3.**
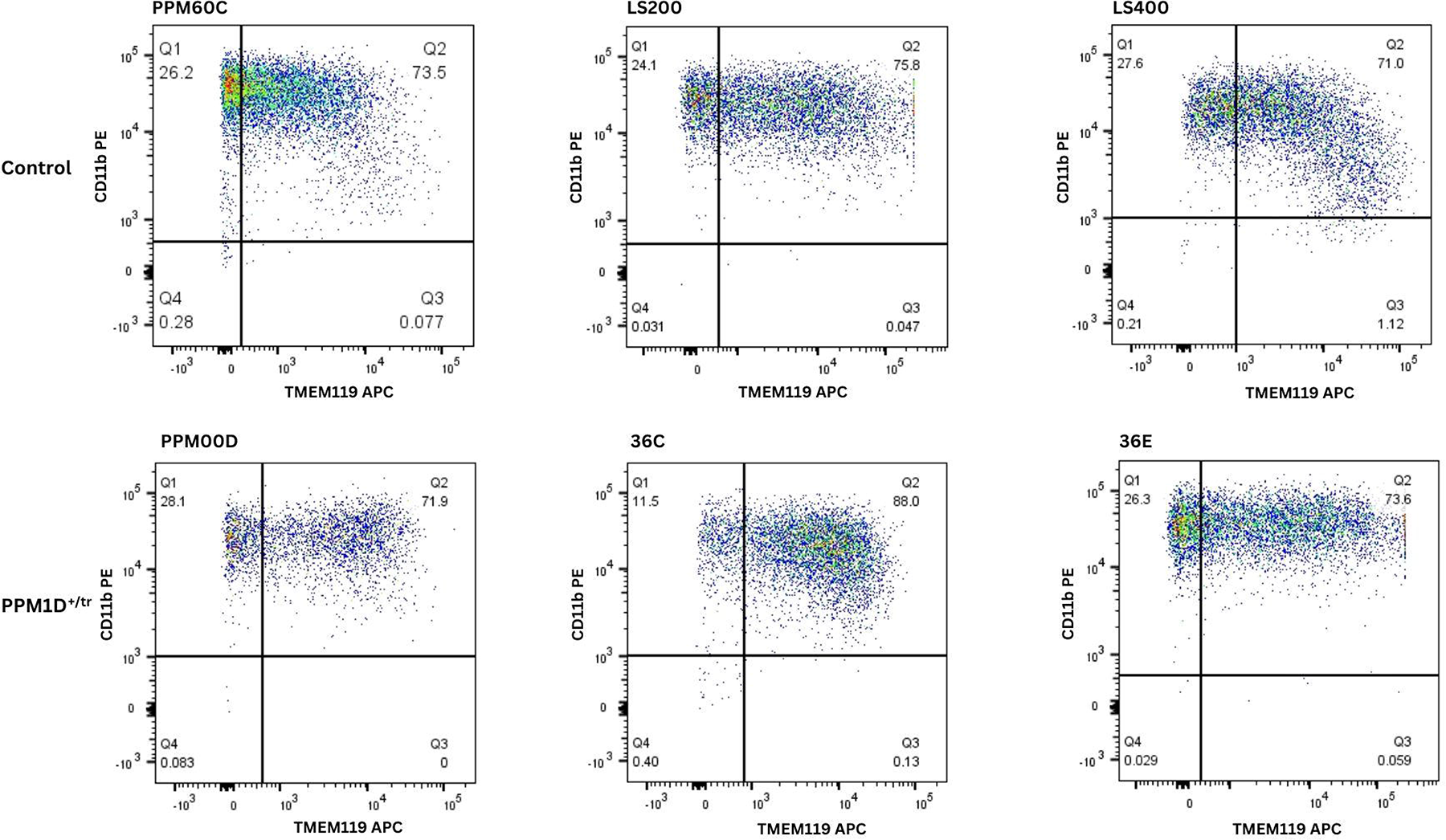
FACS analysis of microglia. Microglia were developed from three control iPSC lines (PPM60C, LS200, and LS400), and three patient lines (PPMOOD, 36C, 36E). PPM6OC is the typically developing sibling control of PPMOOD, and LS200 is the isogenic control for 36C and 36E. LS400 is another typically developing control. Cells were sorted using conjugated antibodies against the microglia markers TMEM119 and CD11b, the latter of which also binds to macrophages. Microglia are positive for both double-positive cells.

**Table 4.**
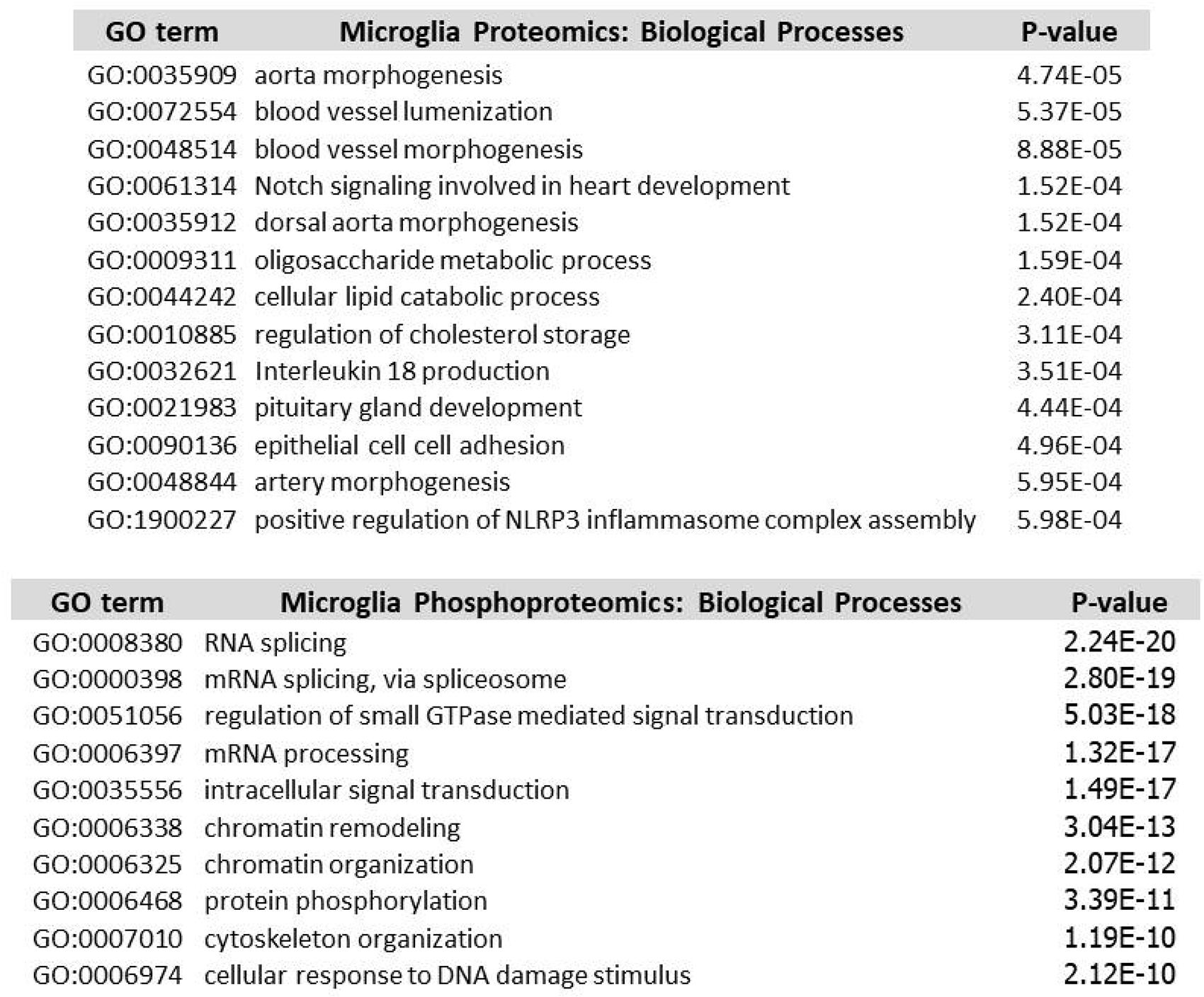
Gene Ontology (GO) analysis, microglia. All differentially expressed proteins and phosphoproteins were used to determine GO terms. Only the most significant terms are shown. For the complete list, see **Additional file 5: Table S3.**

Also consistent with a neuroinflammatory phenomenon are the GO terms of enriched DEPs showing an effect on the production and positive regulation of NLRP3 inflammasome complex assembly. IL-18 is a proinflammatory cytokine produced, along with IL-1β, as a result of NLRP3 inflammasome activation (79–81). However, the level of significance for this GO term is modest.

The top pathway for upregulated proteins was lysosome, but similar to the KEGG analysis of neurons, among the top pathways for both up and downregulated proteins are neurodegenerative disorders. The lysosome pathway could indicate a predilection for disruption of autophagy, a process linked to neurodegenerative and neurodevelopmental disorders (**Table 5**) (82–85). Interestingly, the most upregulated DEPs proteins in *PPM1D*^+/tr^ microglia are several regulators of ubiquitin signaling and innate immune pathways (**Table 6**). These included, CDC34, a Cullin-Ring E2 ubiquitin-conjugating enzyme, GBP5, a member of the GTPase subfamily induced by interferon-gamma (IFN-γ), CDBP2, a CD2 antigen cytoplasmic tail-binding protein that regulates T-cell activation and IL-2 production, KLHDC4, a member of the Kelch-like proteins that act as substrate adaptors for Cullin 3 ubiquitin ligases, PELI1, an E3 ubiquitin protein ligase pellino homolog that regulates NLRP3-induced caspase-1 activation and IL-1β maturation, ZNFX1, which functions as a dsRNA sensor and regulator of antiviral responses, and TEP1, a telomerase protein component that can influence innate immune responses through cGAS/STING (cyclic GMP-AMP synthase-stimulator of interferon genes) activation, a cytosolic DNA sensor (86–92).

**Table 5.** KEGG analysis, microglia. Shows lists of the most significant KEGG pathways for up and downregulated proteins and phosphoproteins in *PPM1D*^+/tr^ microglia. See **Additional file 5: Table S3** for complete lists.

| KEGG pathway up-regulated proteins microglia | p-value | KEGG pathway down-regulated proteins microglia | p-value |
| --- | --- | --- | --- |
| Lysosome | 6.94E-07 | Amyotrophic lateral sclerosis | 1.16E-08 |
| Parkinson disease | 4.53E-06 | Huntington disease | 8.41E-07 |
| Prion disease | 1.77E-05 | Phagosome | 1.35E-06 |
| RNA transport | 6.67E-05 | Salmonella infection | 1.71E-05 |
| Citrate cycle (TCA cycle) | 8.17E-05 | Prion disease | 6.06E-05 |
| Alzheimer disease | 1.52E-04 | Pathways of neurodegeneration | 6.77E-05 |
| Cholesterol metabolism | 2.22E-04 | Proteasome | 1.33E-04 |
| Aminoacyl-tRNA biosynthesis | 2.23E-04 | Pathogenic Escherichia coli infection | 1.33E-04 |
| Glycolysis / Gluconeogenesis | 2.48E-04 | Vibrio cholerae infection | 2.28E-04 |
| Phagosome | 4.26E-04 | RNA transport | 2.49E-04 |

| KEGG pathway up-regulated phosphoproteins microglia | p-value | KEGG pathway down-regulated phosphoproteins microglia | p-value |
| --- | --- | --- | --- |
| Spliceosome | 1.47E-07 | Spliceosome | 5.28E-08 |
| Yersinia infection | 3.92E-05 | Viral life cycle - HIV-1 | 7.29E-07 |
| Rap1 signaling pathway | 7.68E-05 | Regulation of actin cytoskeleton | 8.36E-07 |
| Shigellosis | 1.60E-04 | Thyroid hormone signaling pathway | 1.97E-06 |
| Insulin signaling pathway | 7.70E-03 | Platelet activation | 1.05E-05 |
| NOD-like receptor signaling pathway | 1.79E-02 | Fc gamma R-mediated phagocytosis | 2.77E-05 |
| Transcriptional misregulation in cancer | 2.41E-02 | Endocytosis | 4.57E-05 |
| Adherens junction | 3.31E-02 | Renal cell carcinoma | 5.48E-05 |
| Fc gamma R-mediated phagocytosis | 3.56E-02 | ErbB signaling pathway | 5.51E-05 |
| Platelet activation | 3.71E-02 | Insulin signaling pathway | 5.53E-05 |

**Table 6.**
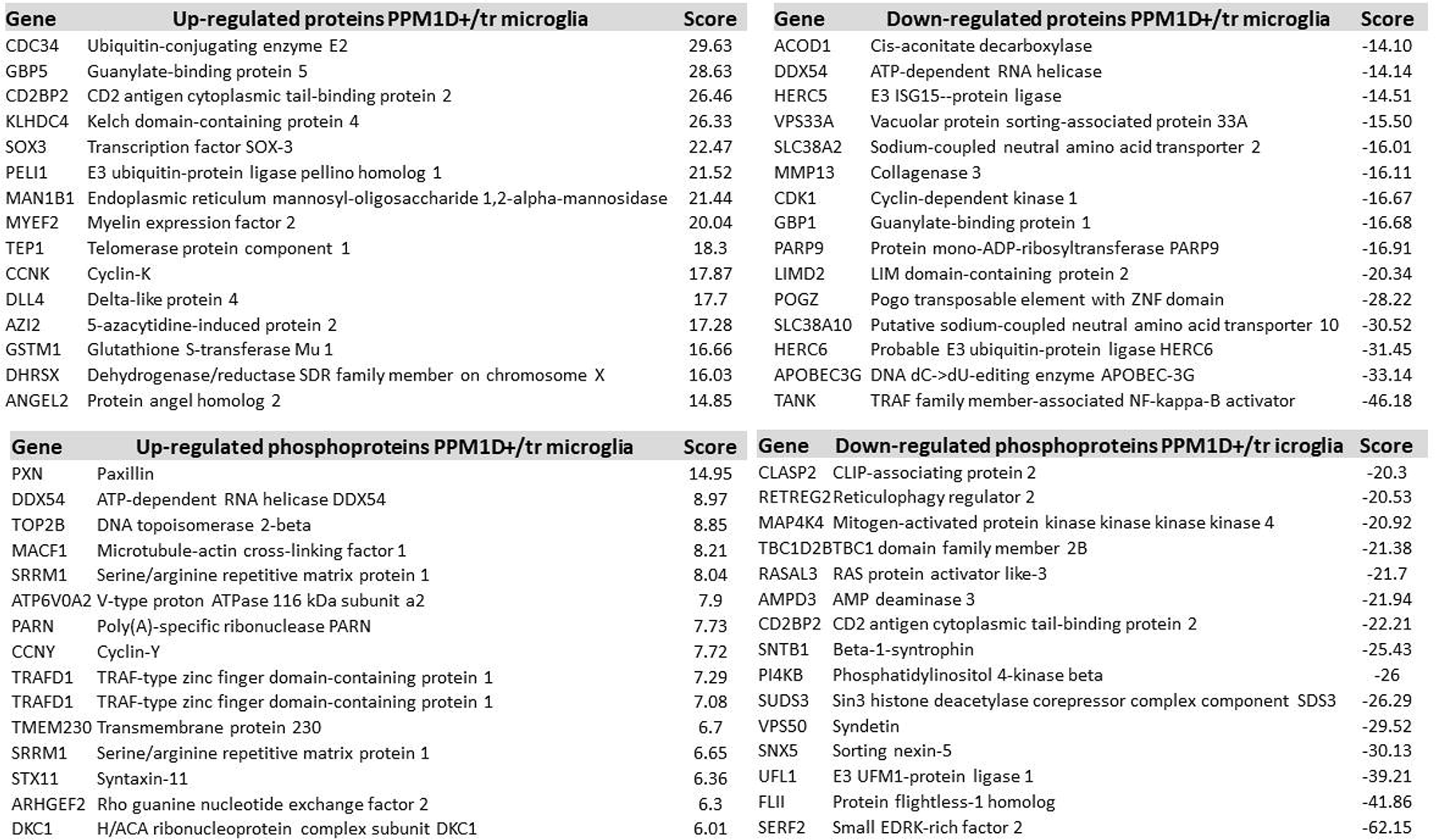
Differentially expressed proteins and phosphoproteins, uninduced microglia. Differentially expressed up and down-regulated proteins and phosphoproteins in baseline, untreated microglia.

Many of the most downregulated proteins are also involved in innate immunity including TANK, which activates NF-κB and cGAS-STING signaling, APOBEC3G, a cytidine deaminase involved in anti-viral innate immunity, and HERC6, an E3 ligase for ISG15 that regulates ISGylation, a post-translational modification induced by interferon that has ubiquitin-like, protein modifying effects (93–103).

Another key downregulated protein is POGZ, a chromatin regulator that also promotes homology-directed DNA repair (104). LOF mutations are commonly found in ASD and NDD (105,106). POGZ binds to ADNP, and their deficiency in mice induces significant upregulation of genes enriched in neuroinflammation and altered microglial and glutamatergic neuronal function (105–108). LOF mutations in ADNP have been found in NDD and ASD (including regressive autism; see discussion) (109–111). Neither POGZ nor ADNP is significantly differentially expressed in glutamatergic neurons. These findings suggest that decreased POGZ expression in microglia is playing a role in the neurodevelopmental features of JdVS.

### Phosphoproteomics: Microglia

The phosphoproteomics analysis detected 3458 phosphosites of which only 4 showed a significant increase in the *PPM1D*^+/tr^ microglia, while 39 were significantly decreased (**Additional file 6: Table S4; Figure 2**). The top GO terms for differentially phosphorylated proteins were related to RNA splicing, regulation of small GTPase signaling, chromatin remodeling, protein phosphorylation, cytoskeleton organization, and cellular response to DNA damage stimulus, and the top KEGG pathway was spliceosome, similar to the neuronal proteomics and phosphoproteomics studies (**Table 4**; **Table 5**). The top upregulated phosphorylated proteins in *PPM1D*^+/tr^ microglia were in PXN, DDX54, TOP2B, MACF1, CCNY, and SRRM1 (**Table 6**). Remarkably, this overlaps with the finding that SRRM1 was the top upregulated, as well as the top downregulated phosphorylated protein in *PPM1D*^+/tr^ neurons, as described above, providing additional support for the idea that PPM1D truncating mutations disrupt SRRM1 function. An increase in phosphorylation at S429 in both neurons and microglia was detected. This is predicted to be a substrate for the PIM family of kinases, which promote tumorigenesis and immune escape by HIV (112,113).

Two phosphosites in TRAFD1 were among the top 10 DEPPs. TRAFD1 is a transcription factor that acts as a negative feedback regulator of the innate immune system to control excessive immune responses (114,115). A similar occurrence in microglia could perhaps be relevant to the acute neuropsychiatric decompensation that occurs in some JdVS patients.

The most downregulated DEPPs were found in SERF2, FLI1, UFL1, SNX5, and VPS50. SERF2 is small EDRK-rich factor 2 that modifies amyloid fiber assembly and promotes protein misfolding (116). FLI1 is a member of the ETS transcription factor family that is disrupted in Ewing Sarcoma and acute myelogenous leukemia (117). And UFL1 (UFM1-protein ligase 1; Ubiquitin-like modifier 1 ligating enzyme 1) is a regulator of UFM1 conjugation (UFMylation), a ubiquitin-like modification that plays a key role in maintaining cell homeostasis under cellular stress, including DDR (118–120). SNX5 is a component of an autophagosomal complex and VPS50 is an endosome-recycling protein (121,122).

In summary, the microglia proteomics and phosphoproteomics analyses suggest that reduced expression of POGZ is a candidate for the cognitive and behavioral aspects of JdVS, and that truncated PPM1D could disturb innate immune responses in the brain through altered regulation of ubiquitin signaling, DDR, splicing, altered expression or function of key genes, and perhaps BBB permeability.

### Proteomics Analysis of lipopolysaccharide (LPS)-activated microglia

To test the hypothesis that PPM1D^+/tr^ microglia have an altered response to an innate immune system challenge, the effect of LPS was analyzed by proteomics. One hundred and fifty-eight proteins were upregulated in the PPM1D+/tr samples, and 254 were downregulated. (**Additional file 7: Table S5; Figure 2**). The top GO term was negative regulation of interleukin-6 (IL-6), a proinflammatory cytokine implicated in neuroinflammation, maternal immune activation, ASD, schizophrenia, and depression (**Table 7**) (123–126). However, the p-value was modest. KEGG analysis showed that the top pathways for upregulated proteins were lysosome, metabolic pathways, and several neurodegenerative disorders, similar to the findings in uninduced microglia and glutamatergic neurons (**Table 8**). Downregulated proteins were enriched for spliceosome, nucleocytoplasmic transport, and ALS, overlapping with other proteomics findings. Proteins involved in the response to several infectious diseases were also detected in the KEGG analysis.

**Table 7.** Gene Ontology (GO) analysis, LPS induced microglia. All differentially expressed proteins and phosphoproteins were used to determine GO terms. Only the most significant terms are shown. For the complete list, see **Additional file 7: Table S5**.

| **GO term** | **LPS Microglia Proteomics: Biological Processes** | **P-value** |
| --- | --- | --- |
| GO:0032715 | negative regulation of interleukin-6 production | 3.23E-05 |
| GO:0072310 | glomerular epithelial cell development | 3.44E-05 |
| GO:0072015 | glomerular visceral epithelial cell development | 3.44E-05 |
| GO:0034446 | substrate adhesion-dependent cell spreading | 1.08E-04 |
| GO:0015850 | organic hydroxy compound transport | 1.39E-04 |
| GO:0031589 | cell-substrate adhesion | 1.63E-04 |
| GO:0072359 | circulatory system development | 1.90E-04 |
| GO:2000351 | regulation of endothelial cell apoptotic process | 2.08E-04 |
| GO:0003014 | renal system process | 2.51E-04 |
| GO:0006003 | fructose 2,6-bisphosphate metabolic process | 3.31E-04 |

| **GO term** | **LPS Microglia Phosphoproteomics: Biological Processes** | **P-value** |
| --- | --- | --- |
| GO:0008380 | RNA splicing | 2.32E-20 |
| GO:0000398 | mRNA splicing, via spliceosome | 2.89E-19 |
| GO:0051056 | regulation of small GTPase mediated signal transduction | 5.17E-18 |
| GO:0006397 | mRNA processing | 1.37E-17 |
| **GO:0035556** | intracellular signal transduction | 1.56E-17 |
| **GO:0006338** | chromatin remodeling | 3.11E-13 |
| **GO:0006325** | chromatin organization | 2.14E-12 |
| **GO:0006468** | protein phosphorylation | 3.52E-11 |
| **GO:0007010** | cytoskeleton organization | 1.22E-10 |
| **GO:0006974** | cellular response to DNA damage stimulus | 2.18E-10 |
| **GO:0045944** | positive regulation of transcription from RNA polymerase II promoter | 2.86E-10 |
| **GO:0051301** | cell division | 4.59E-09 |
| **GO:0030036** | actin cytoskeleton organization | 9.04E-09 |
| **GO:0007165** | signal transduction | 1.01E-08 |
| **GO:0090630** | activation of GTPase activity | 2.75E-08 |

**Table 8.** KEGG analysis, LPS treated microglia. Differentially expressed up and down regulated proteins and phosphoproteins in baseline, untreated microglia. **Additional file 7: Table S5.**

| KEGG pathway up-regulated proteins LPS microglia | p-value | KEGG pathway down-regulated proteins LPS microglia | p-value |
| --- | --- | --- | --- |
| Lysosome | 2.38E-22 | Spliceosome | 6.00E-15 |
| Metabolic pathways | 1.22E-13 | Nucleocytoplasmic transport | 2.12E-13 |
| Huntington disease | 6.41E-12 | Amyotrophic lateral sclerosis | 1.68E-12 |
| Parkinson disease | 1.14E-11 | Salmonella infection | 1.86E-12 |
| Prion disease | 5.47E-11 | Ribosome | 1.15E-10 |
| Amyotrophic lateral sclerosis | 5.06E-10 | Protein processing in endoplasmic reticulum | 2.03E-10 |
| Oxidative phosphorylation | 1.25E-09 | Epstein-Barr virus infection | 2.74E-10 |
| Chemical carcinogenesis - reactive oxygen species | 4.44E-08 | Carbon metabolism | 1.08E-09 |
| Pathways of neurodegeneration - multiple diseases | 5.90E-08 | Shigellosis | 1.67E-09 |
| Neutrophil extracellular trap formation | 6.17E-08 | Leishmaniasis | 2.50E-08 |
| Proteasome | 1.43E-07 | Metabolic pathways | 2.64E-08 |
|  |  | Coronavirus disease - COVID-19 | 3.92E-08 |
|  |  | Influenza A | 7.27E-08 |

| KEGG pathway up-regulated phosphoproteins LPS microglia | p-value | KEGG pathway down-regulated phosphoproteins LPS microglia | p-value |
| --- | --- | --- | --- |
| Rap1 signaling pathway | 6.29E-06 | Spliceosome | 2.07E-09 |
| Yersinia infection | 7.87E-06 | Regulation of actin cytoskeleton | 7.01E-07 |
| Platelet activation | 8.81E-06 | Viral life cycle - HIV-1 | 7.27E-05 |
| Regulation of actin cytoskeleton | 7.58E-05 | Thyroid hormone signaling pathway | 9.62E-05 |
| Fc gamma R-mediated phagocytosis | 1.78E-04 | ErbB signaling pathway | 1.37E-04 |
| Adherens junction | 2.10E-04 | Shigellosis | 1.59E-04 |
| Shigellosis | 2.24E-04 | Insulin signaling pathway | 1.78E-04 |
| Pathogenic Escherichia coli infection | 2.59E-04 | Rap1 signaling pathway | 3.22E-04 |
| Thyroid hormone signaling pathway | 4.00E-04 | Leukocyte transendothelial migration | 4.17E-04 |
| Chemokine signaling pathway | 5.52E-04 | Yersinia infection | 5.00E-04 |

Examination of individual DEPs showed striking patterns consistent with innate immune dysregulation, in particular, ubiquitin signaling (**Table 9**). The top upregulated protein in PPM1D^+/tr^ microglia was UHRF1BP1 (UHRF1-binding protein 1), which binds to UHRF1, a RING-finger E3 ubiquitin ligase, a regulator of Treg cell proliferation (127,128). Non synonymous variants have been found in systemic lupus erythematosus (129). However, the effects of UHRF1BP1 and UHRF1 on microglia are not known. GBA1 catalyzes the cleavage of glycosphingolipids–glucosylceramide and glucosyl sphingosine. Genetic variants are known risk factors for PD and Lewy Body Dementia, and biallelic LOF variants cause Gaucher disease (130–134). RAB11A is a member of the RAS family of small GTPases and is a regulator of toll receptor trafficking (134,135).

**Table 9.**
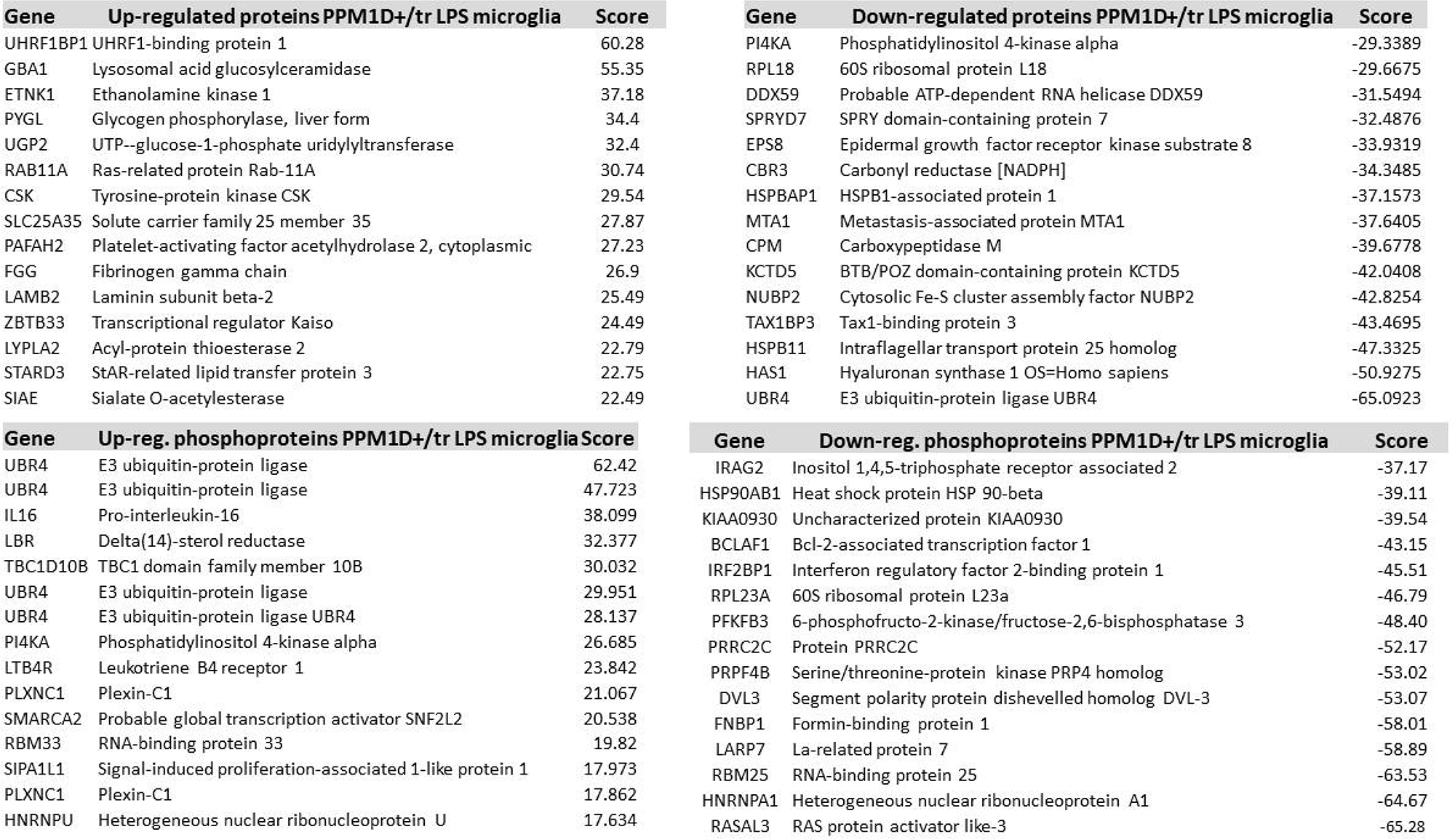
Differentially expressed proteins and phosphoproteins, LPS-treated microglia. Shows the top up and downregulated proteins and phosphoproteins in baseline, untreated microglia. See **Additional file 7: Table S5 and Additional file 8: Table S6** for all proteins and phosphoproteins.

The most downregulated DEP was the ubiquitin protein ligase E3 component n-recognin 4 protein, UBR4, which regulates oxidative stress by promoting K27-linked-ubiquitylation of N-terminal oxidized cysteines leading to proteasomal degradation (136). It’s also a regulator of interferon signaling (137,138) and the proteasomal degradation of PINK1, which is involved in the pathogenesis of PD (see discussion) (138–140). *UBR4* variants have been linked to early onset dementia (140). Other top-downregulated proteins include HAS1, a regulator of the extracellular matrix that is induced by LPS and KCTD5, a BTB/POZ domain-containing protein that functions as substrate-specific adaptor for Cullin3-based E3 ligases (141–143).

### Phosphoproteomics LPS treated microglia

Phosphoproteomics was carried out on the same samples used in the proteomics analysis. A total of 3,458 phosphosites were detected; 42 showed a significant increase in the LPS treated *PPM1D^+/tr^* microglia, and 182 showed a significant decrease (**Additional file 8: Table S6; Figure 2**). Strikingly, four of the top seven phosphosites that increased in the LPS-stimulated *PPM1D*^+/tr^ microglia were in UBR4, although neither of the sites is a canonical PPM1D target motif (**Table 9**). The top phosphosite is at S2718, which is phosphorylated in UV irradiated cells, consistent with an effect of PPM1D or its substrates on DDR (144). This and the other top UBR4 phosphosites have an SS motif. As noted above, UBR4 is also the most downregulated DEP in LPS-treated microglia, suggesting that UBR4 phosphorylation is inversely correlated with UBR4 protein levels. There are also two enriched phosphosites in PLXNC1, a member of the plexin family of transmembrane receptors for semaphorins, which are involved in brain development and immune responses (145). IL16 had the most differentially increased phosphosite after UBR4. It is a CD4+ immune cell-specific chemoattractant cytokine that has been implicated in multiple sclerosis (146,147).

The most downregulated phosphosite was in RASAL3, a RasGAP that is highly expressed in neutrophils. Deficiency enhances immune activation in acute inflammatory conditions (148). GO analysis of all differential phosphosites showed enrichment of phosphoproteins involved in RNA splicing, regulation of small GTPase-mediated signal transduction, chromatin remodeling, cytoskeleton organization, and cellular response to DNA damage stimulus (**Table 7**). These findings overlap with the uninduced microglia phosphoproteomics analysis. Alterations in microglia’s cytoskeleton function could potentially cause problems with their migration or phagocytic potential.

The top KEGG pathways for phosphosites that increased in the *PPM1D*^+/tr^ microglia were Rap1 signaling, Yesinia infection, platelet activation and regulation of cytoskeleton, while the terms spliceosome, regulation of actin cytoskeleton, viral life cycle (HIV-1) and thyroid hormone signaling pathway were pathways enriched with phosphosites that decreased in *PPM1D*^+/tr^ microglia (**Table 8**). A modest enrichment of phosphosites involved in ErbB signaling was also seen, which by itself seems relatively minor. However, in view of the enrichment of phosphosites in this pathway in neurons and uninduced microglia, as well as LPS-treated cells, the findings suggest that *PPM1D* truncating mutations can disrupt ErbB signaling. PPM1D expression has been found to affect breast cancer growth (149,150). In the brain, ErbB signaling plays a role in synaptic plasticity and has been implicated in NDD (151–154).

Overall, the findings show that immune stimulation of PPM1D^+/tr^ microglia results in altered expression of proteins involved in ubiquitin signaling, in particular, UBR4, actin cytoskeleton, RNA splicing, chromatin structure, and innate immune regulation, the latter of which could play a role in the decompensation some JdVS patients experience following infectious and non infectious stressors.

In summary, the three cell types in which proteomics and phosphoproteomics were carried out (glutamatergic neurons, uninduced microglia, and LPS-stimulated microglia) showed several overlapping pathways: splicing, ubiquitin ligase expression, neurodegenerative disorders, chromatin organization, cytoskeleton dynamics, and ErbB signaling (**Figure 4**).

**Figure 4.**
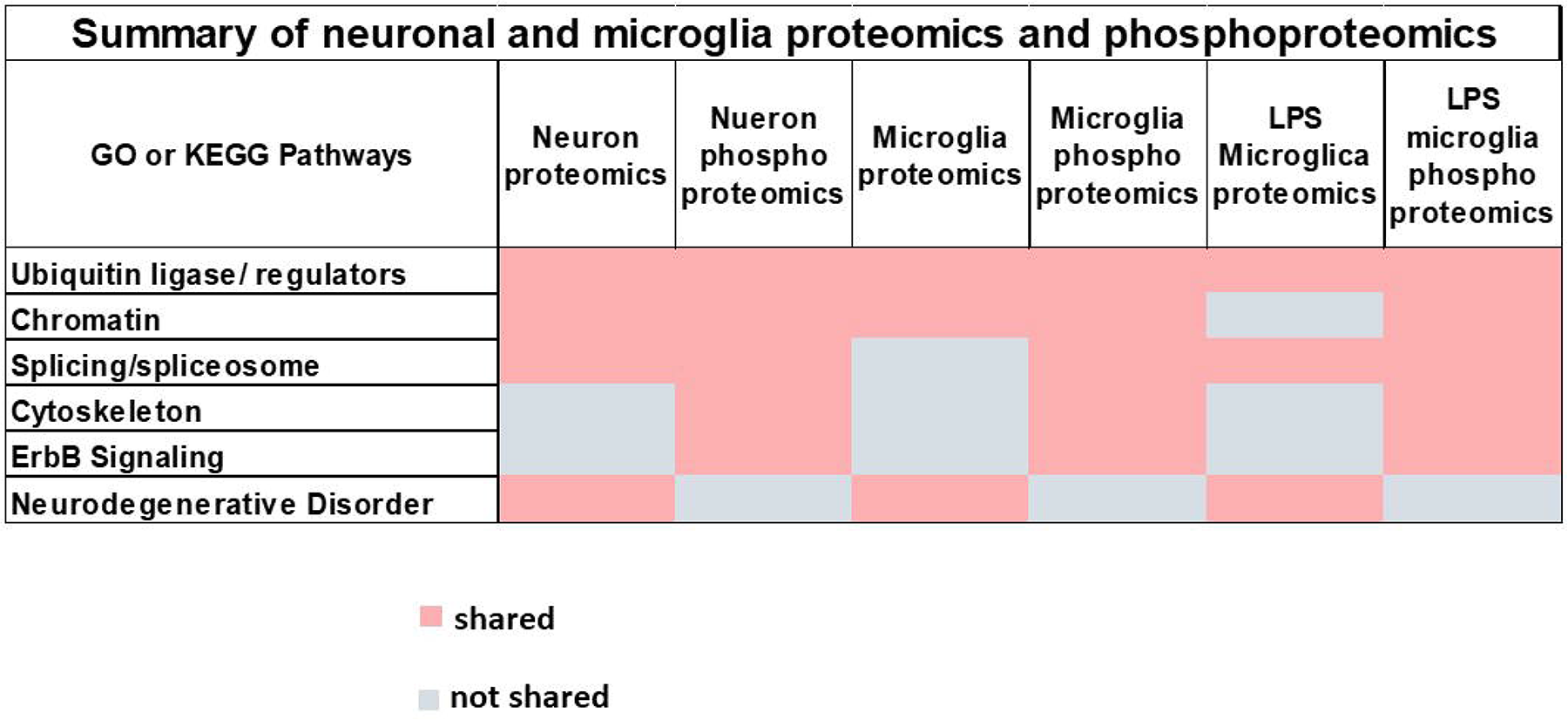
Summary of neuronal and microglia proteomics and phosphoproteomics. Six pathways (center of each block, light grey) were found by either GO or KEGG (up or downregulated) that were shared in three or more of the six different conditions analyzed in this study; proteomics and phosphoproteomics on glutamatergic neurons, untreated microglia, and LPS induced microglia (shared dark grey block; not shared, white block)

### Functional analysis of glutamatergic neurons and microglia

Most of the differentially expressed phosphosites in neurons and microglia were not at canonical SQ or TQ motifs recognized by PPM1D, so the GOF effect of truncated PPM1D could not be unequivocally validated using the phosphoproteomics data we obtained. Consequently, we examined a known neuronal PPM1D target, CaMKII T287 to confirm a GOF effect in neuronal cells. As shown in **Figure 5**, there was a statistically significant, 2-fold decrease in the relative expression of phospho-CAMKII in *PPM1D*^+/tr^ neuronal cells consistent with a GOF effect. Also shown in the figure, is a validation of the GO and KEGG findings that cytoskeleton function is disrupted in *PPM1D*^+/tr^ glutamatergic neurons; a significant decrease in neurite outgrowth was found. Cytoskeleton function is critical for neurite outgrowth and synapse development, and many ASD and NDD candidate genes have an adverse effect on these processes (155–157).

**Figure 5.**
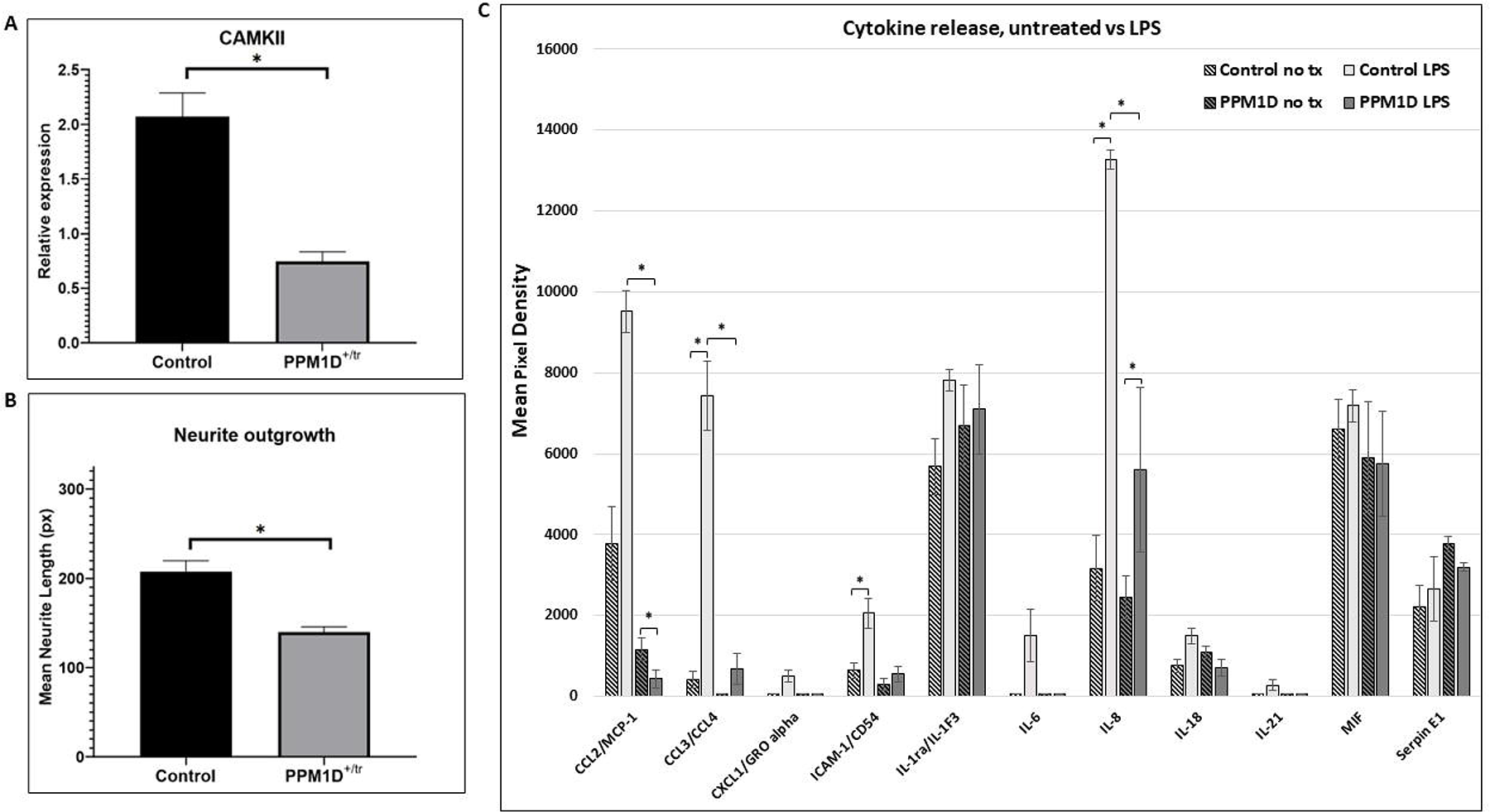
CaMKII, Neurite Outgrowth, Cytokine Release. **A.** CaMKII phosphorylation was analyzed by quantifying Western blot signals for CaMKII, phospho-CaMKII, and cyclophilin as a loading control. The graph in 5A is a plot of the ratio of the normalized phospho-CaMKII signal (relative to cyclophilin) divided by the normalized CaMKII signal. A total of 4 control and 5 *PPM1D^+/tr^* neuronal samples were analyzed. The graph is the mean for the two groups, +/- SEM. The decrease in CaMKII phosphorylation was highly significant (p= 0.0005, Student’s t-test, two-tailed). **B.** Neurite outgrowth was measured in day 21 glutamatergic neurons as described in the methods section. Tracings from two control and two *PPM1D^+/tr^* day 21 glutamatergic neurons were obtained, blind to genotype, for a total of 400 and 229 neurons analyzed, respectively. The difference between the two was highly significant (mean +/- SEM, 8.9E-07, Student’s t-test, two-tailed). **C.** Cytokine release was assayed using the Proteome Profiler Array Human Cytokine Array, as described in the methods. A total of 4 control and 3 *PPM1D^+/tr^*microglia samples were analyzed (the same samples that were analyzed in the proteomics and phosphoproteomics analyses plus an additional control that was not analyzed by proteomics. Culture supernatants were harvested after 24 hours of LPS treatment. Untreated (no tx) cells from the same differentiation were harvested simultaneously. The data are the means of 4 vs 3, with each spot on the array measured in duplicate. A Student’s t-test was used to calculate statistical significance. Those at p < 0.05 are indicated by asterisks: CCL2, untreated control LPS vs *PPM1D*^+/tr^ LPS (p=0.01); untreated *PPM1D*^+/tr^ vs *PPM1D*^+/tr^ LPS (p=0.0009) (note CCL2 control vs control LPS had a p-value of 0.065); CCL3/CCL4, untreated control vs control LPS (p=0.02); CCL3/CCL4, control LPS vs *PPM1D*^+/tr^ LPS (p=0.02); ICAM-1 untreated control vs control LPS (p=0.05); IL-8, untreated control vs control LPS (p=0.02), control LPS vs *PPM1D*^+/tr^ LPS (p=0.05);

Finally, we measured the concentration of cytokines and chemokines in the supernatant following LPS treatment to assess microglia function following an innate immune challenge. The top GO term for LPS-treated *PPM1D*^+/tr^ microglia was negative regulation of IL-6. This was based on DEPs that directly or indirectly affect IL-6 signaling (ZC3H12A, GBA, TNFAIP3, GAS6), rather than IL-6 levels per se, which was not detected in the proteomics analysis. As seen in **Figure 5C**, IL-6 was induced in LPS-treated control microglia, but not in the *PPM1D*^+/tr^ cells as predicted. However, because of a large standard error and the small sample size, the induction was not statistically significant. In addition, we detected a slight increase in baseline IL-18 levels in *PPM1D*^+/tr^ microglia compared with the controls, as predicted from the GO analysis, but that difference was also not statistically significant. There was also a decrease in IL-18 induction by LPS detected in the *PPM1D*^+/tr^ microglia compared with LPS-treated controls, but the difference fell short of statistical significance (p=0.2). This is consistent with the proteomics data, which showed a statistical trend towards a decrease in IL-18 in the *PPM1D*^+/tr^ LPS-treated microglia (-log2 p-value of 3.79 = 0.07, see **Additional file 7: Table S5**). In addition, there was, somewhat unexpectedly, a significant decrease in the induction of the chemokines CCL2 and CCL3/CCL4 by LPS between the control and *PPM1D*^+/tr^ microglia (p= 0.0009 and 0.02, respectively. In fact, while LPS induced an increase in CCL2 in the control sample that showed a trend towards statistical significance (p=0.065), a significant decrease was detected in the *PPM1D*^+/tr^ cells (p=0.01). The findings suggest that truncated PPM1D causes deregulation of cytokine release.

## Discussion

Although the sample size in this study was small, a number of interesting findings emerged that could explain the clinical features seen in JdVS. One is the significant decrease in the expression of the chromatin regulator and high-confidence ASD candidate gene *POGZ* in *PPM1D*^+/tr^ microglia. This is consistent with the finding that *Pogz* KO mice show an upregulation of genes enriched in neuroinflammation and an increase in microglia phagocytosis in the prefrontal cortex (105). Similar and overlapping findings were found in *Adnp* KO mice. *ADNP* is another high-confidence ASD gene that forms a nuclear complex with POGZ and was also decreased in *PPM1D*^+/tr^ microglia (108). Given the findings in mouse *Pogz* KO models, the significant decrease of POGZ in *PPM1D*^+/tr^ suggests that microglia are playing a direct role in the neurodevelopmental aspects of JdVS. However, considering the neuronal proteomics analysis, neurons are likely also playing a causal role, as expected of a gene like *PPM1D* that is expressed ubiquitously throughout the CNS in neurons and non-neuronal cells. Considering that *PPM1D* is expressed at much higher levels in the cerebellum compared to other brain regions, as are both *POGZ* and *ADNP*, it will be particularly important to carry out the studies reported here in cerebellar organoids derived from our iPSC lines, or in mouse models. The cerebellum is now known to play a role in cognitive function and the development of ASD and NDD, in addition to its well-established effects on locomotor function and coordination (158–160).

Another interesting aspect of this study is related to the clinical observation that a small proportion of JdVS patients have acute neuropsychiatric decompensation, a phenomenon that has been described in genetic subgroups of NDD and ASD (109,161–164). One of the patients presented with symptoms consistent with PANS, as we previously reported (15), and two others with acute and subacute behavioral and motor regression following infection and noninfectious stressors (unpublished observations). PANS is an autoinflammatory/autoimmune disorder induced by group A beta-hemolytic *Streptococcus* and other infectious microbes, and behavioral regression in ASD has been hypothesized to have immune-based or infection-triggered underlying pathogenesis in some cases (162,165–167). Interestingly, *ADNP* is one of 11 ASD associated candidate genes, most commonly found in regressive ASD (109). Our microglia proteomics and phosphoproteomics findings support the idea that *PPM1D* truncating mutations can affect the susceptibility to immune-based decompensation. For example, the microglia GO analysis showed that proteins involved in endothelial function that can affect the BBB are differentially expressed; disruption of the BBB resulting in increased permeability to peripheral inflammatory cytokines, chemokines, complement, and immune cells, has been viewed as a pathogenic feature of neuroinflammatory and neurodegenerative disorders (168–171). Also consistent with a neuroinflammatory vulnerability are the various connections to dysregulated innate immunity we detected in *PPM1D*^+/-^ microglia, primarily through altered expression of ubiquitin ligases and ubiquitin-conjugating enzymes. Most notable is UBR4, which was the most downregulated DEP in LPS-stimulated microglia. UBR4 was also among the most down regulated proteins detected in *PPM1D*^+/tr^ microglia treated with poly I:C and IL-17 (unpublished observations, manuscript in preparation). As noted in the results section, UBR4 regulates oxidative stress and interferon signaling, and the degradation of PINK1, a mitochondrial serine/threonine kinase that recruits the E3 ligase PARKIN (PRKN) to induce mitophagy (136–138,172). Homozygous, LOF mutations in either *PINK1* or *PRKN* are found in early onset, autosomal recessive forms of PD (139,140). Interestingly, we are aware of two cases of acute neuropsychiatric decompensation consistent with PANS who are heterozygous for LOF mutations in *PRKN* (manuscript in preparation). This connection between LPS-stimulated *PPM1D*^+/tr^ microglia and PD suggests a common vulnerability to environmental stressors due to dysfunctional UBR4 signaling, with subsequent adverse effects on proteasomal degradation and mitophagy. A defect in mitophagy homeostasis has been implicated in the pathogenesis of PD, as well as AD (173,174). *UBR4* variants have been found in some families with early-onset dementia (175). The effect of *PPM1D* truncating mutations on mitophagy has not yet been carried out. It should be noted that DEPs aside from UBR4 are connected to PD as seen in the KEGG pathway analyses showing that PD (and other neurodegenerative disorders) is among the top GO and KEGG pathways in both neurons and microglia.

In addition to UBR4, a number of other E3 ubiquitin ligases and their regulators, and ubiquitin conjugating enzymes involved in innate immune responses were among the most differentially expressed proteins in untreated *PPM1D*^+/tr^ microglia (CDC34, KLHDC4, and PELI1; upregulated: HERC6; downregulated), and in LPS stimulated microglia (UHRF1BP1, upregulated), as noted in the results section. In addition, the top-upregulated protein in *PPM1D*^+/tr^ neurons was CUL4B, a component of the CUL4B-Ring E3 ligase. The specific substrates affected by these ubiquitin regulators in microglia and neurons, and how they might be involved in JdVS and neuroinflammation need to be investigated.

An important finding to consider regarding the neuroinflammatory potential of *PPM1D*^+/tr^ microglia is the seemingly paradoxically lack of induction of IL-6 following LPS stimulation and the GO analysis that showed an enrichment of proteins that negatively regulate IL-6. This cytokine is one of the major proinflammatory cytokines implicated in neuroinflammation and maternal immune activation (123,176,177) In fact, in addition to IL-6, there was a generalized blunting of LPS-mediated cytokine induction in *PPM1D*^+/tr^ microglia (**Figure 5C**). This was particularly the case for CCL2 and CCL3/CCL4, which showed significant decreases compared with LPS-induced control microglia. A blunted induction of ICAM-1 and IL-8 was also detected, but the control vs PPM1D^+/tr^ difference was only significant for the latter (p=0.05). This suggests that TLR4 signaling is attenuated by truncated PPM1D. These findings need to be validated in a larger iPSC dataset, which is currently being carried out, and animal models. Reduced expression of CCL2 and CCL3/CCL4, which are potent monocyte attractants, in response to LPS activation, could affect the recruitment of transmigrating monocytes, a population of peripheral monocytes that can cross the BBB (178,179). Considering the dichotomy of monocytes and macrophages into pro- and anti-inflammatory subtypes, the effect of the blunted activation of CCL2 and CCL3/CCL4 on a neuroinflammatory process is difficult to predict and would need to be evaluated in an animal model. Furthermore, if the connection between *PPM1D* GOF variants and neuroinflammation is confirmed, then the idea that cytokines and chemokines affecting microglia and brain function are mediators may be too limiting. In the case of truncated PPM1D, a disturbance in microglia homeostasis and function may be at play, rather than an excessive release of proinflammatory cytokines and chemokines.

It is important to note that the iPSC lines used in these studies were not made from subjects with acute psychiatric decompensation, in either the JdVS subject we used or, especially, the CRISPR lines, which were made from a typically developing control. Thus, while connections to a neuroinflammatory process in this study are based on differences between control and *PPM1D* truncating mutations, they are occurring in the context of cellular stress conditions inherent in growing cultured cells in artificial media in an incubator. Therefore, replication in an animal model is essential.

Finally, it is important to consider the microglia findings in the context of the increased infections that occur in some JdVS children. In the original report of 14 cases, and a subsequent follow up of 33 cases, recurrent infections, in particular otitis media, were reported in approximately half (1,4). Although this has not been evaluated systematically, the increase in infections seen in some children has been sufficiently severe to warrant evaluations for immunodeficiencies by their physicians, which have been non-diagnostic so far. Considering the physiological and functional overlap between microglia and peripheral monocytes/macrophages, the differential expression of proteins involved in immune responses that were found in our proteomics analysis, as well as the blunted effect on cytokine and chemokine release could be having a similar effect in the periphery, reducing the effectiveness of an innate immune response to infections.

### Limitations

The small sample size was a major limitation. Another is extrapolating data related to neuroinflammation using an in vitro microglia differentiation and cell culture system that is probably inducing cellular stress.

## Conclusion

In summary, our findings show plausible mechanisms for the neurodevelopmental and cognitive features of JdVS, as well as the increased risk of neuroinflammatory-mediated decompensation, and perhaps the increased rate of infections seen in patients. The mechanistic links we identified to regression in NDD are also significant. Our findings provide additional support for the idea that a subgroup of NDD and ASD cases can experience neuropsychiatric decompensation caused by dysregulated innate immunity that is potentially treatable with immune modulators, as suggested by other investigators (162,180). In addition, the molecular connection to PD found with UBR4 and other DEPs (e.g., NUCKS1, GBA1) could also be significant in that unrecognized and under-treated neuroinflammatory processes could pose a future risk of PD and other neurodegenerative disorders. Thus, our analysis of JdVS, a rare disease with fewer than 100 reported cases, could be informative for disorders that have greater public health significance.

## Supporting information

expanded methods

microglia in suspension

neuron proteomics

neuron phosphoproteomics

microglia proteomics

microglia phosphoproteomics

LPS treated mcroglia proteomics

LPS treated microglia phosphoproteomics

## Abbreviations

JdVS: (Jansen de Vries Syndrome)
iPSC: (induced pluripotent stem cell)
LOF: (loss-of-function)
GOF: (gain of function)
LPS: (lipopolysaccharide)
NDD: (neurodevelopmental disorder)
ADHD: (attention deficit hyperactivity disorder)
ASD: (autism spectrum disorder)
DDR: (DNA damage response)
OCD: (obsessive-compulsive disorder)
KO: (knockout)
hSC: (hematopoietic stem cells)
FACS: (Fluorescence-activated cell sorting)
PANS: (pediatric acute-onset neuropsychiatric syndrome)
GO: (Gene Ontology)
KEGG: (Kyoto Encyclopedia of Genes and Genomes)
BBB: (brain blood barrier)
ALS: (Amyotrophic Lateral Sclerosis)
HD: (Huntington Disease)
PD: (Parkinson Disease)
AD: (Alzheimer Disease)
IFN-γ: (interferon-gamma)
cGAS/STING: (cyclic GMP-AMP synthase-stimulator of interferon genes)

## Declarations

### Ethics approval and consent to participate

Informed consent was obtained by the corresponding author under an Albert Einstein College of Medicine, IRB-approved protocol.

### Consent for publication

Consent for publication was obtained from participants and parents

### Availability of data and materials

Proteomics and phosphoproteomics data were deposited at xxxxxxxxxx ().

### Authors’ contributions

JTA (proteomics and phosphoproteomics, bioinformatics)

EP (prepared iPSCs, differentiated neurons and microglia, proofread manuscript, FACS)

HD (Western blots)

RNB (Western blots)

JB (Neurite outgrowth)

JZ (FACS)

SS (proteomics and phosphoproteomics, bioinformatics)

HML (conceived and designed experiment, analyzed data, wrote manuscript)

## Acknowledgments

We want to thank the Jansen de Vries Syndrome Foundation for their work and the families who provided samples for iPSC development. We also want to thank the Human Pluripotent Stem Cell Core (Director Eric Bouhassira, and Zi Yan) at the Albert Einsein College of Medicine for preparing and characterizing iPSC lines.

## Additional Files

**Additional file 1: Expanded Methods**. This file contains a detailed description of the methods used in this study.

**Additional file 2: Fig. S1.** Microscopic image of unstained microglia grown in suspension. Two controls and two *PPM1D*^+/tr^ samples are shown. The bar is 200 µMs. Two other samples not available.

**Additional file 3: Table S1. Neuron proteomics, PPM1D^+/tr^ vs controls. Tab 1. Gene Ontology (GO) analysis**. Control sample names on top are shown in pastel; 1 and 4, 2 and 3 are biological duplicates. PPM1D^+/tr^ samples are shown in light green. Samples 1 and 2, as well as 3 and 5, are biological duplicates. Differentially expressed proteins are arranged in descending order based on highest to lowest scores in a comparison of all PPM1D^+/tr^ samples vs all controls. The score is calculated by multiplying Fold Change by the p-value (-log 2 of 4.32 corresponds to p=0.05). **Tabs 2 and 3** (Neuron PPM1D^+/tr^ Up KEGG and Neuron PPM1D+/tr Down KEGG, respectively) shows the KEGG analysis of all proteins that increased and decreased in the *PPM1D*^+/-^ samples.

**Additional file 4: Table S2. Neuron phosphoproteomics, PPM1D^+/tr^ vs controls.** Control sample names on top are shown in pastel. Controls 1 and 3, and 2 and 4 are biological duplicates. PPM1D^+/-^ sample names are shown in green. Samples 2 and 4 are biological duplicates. Tab 1, DEPPs are differentially expressed phosphoproteins arranged in descending order based on highest to lowest scores, as described in the legend for Additional file 3: Table S1. Tab 1 (Phosphosites and GO) contains all phosphorylated proteins in descending order and the GO analysis. Tab 2 (normalize to protein value) show GO analysis and volcano plot. Tabs 3 and 4 are neuron phospho up 500 EnrichR and neuron phospho down 500 EnrichR, respectively, showing KEGG pathway analysis for phosphoproteins that increase or decrease in PPM1D^+/-^ neurons. Additional GO analyses for up and down regulated phosphoproteins are shown, but were not described in the paper in order to avoid redundancy.

**Additional file 5: Table S3. Microglia proteomics, PPM1D^+/tr^ vs controls.** Three control samples and three PPM1D^+/-^ samples were analyzed. The microglia were in their baseline state – no treatment (no tx). Tab 1 (processing) shows the detailed steps of data processing starting from raw abundance to log2 transformation, to normalization and to imputation of missing values. Tab 2 is the differentially expressed protein list calculated as described in the neuron proteomics analysis (see figure legend, Additional file 3: Table S1), along with the GO analysis. Tab 3 is the DAVID KEGG analysis of up regulated proteins, which was not discussed in the paper since it overlapped with other findings. Tabs 4 and 5 are the KEGG pathways for the up and down regulated proteins, respectively, using 500 EnrichR. Tab 6 is the DAVID KEGG analysis of up regulated proteins.

**Additional file 6: Table S4; Figure 2. Microglia phosphoproteomics, PPM1D^+/tr^ vs controls**. Phosphoproteomics was carried out on the same samples described in Additional file 5: Table S3. Tab 1 (PPM1D+tr untreated vs Control) contains the differentially expressed phosphoprotein list, along with the GO analysis. Tabs 2 and 3 (PPM1D+tr untreated Up KEGG and PPM1D+tr untreated Down KEGG) show KEGG pathways for up and downregulated phosphosites, respectively.

**Additional file 7: Table S5. LPS treated microglia proteomics, PPM1D^+/tr^ vs controls.** Microglia derived from the same iPSC lines as described in the legend of Additional file 5: Table S3 were treated with LPS (see main text). Tab 1 (processing), as described in Additional file 5: Table S3. Tab 2 (microglia proteomics LPS) shows differentially expressed protein arranged by total score in descending order (*PPM1D*^+/tr^ vs controls) and GO analysis. KEGG pathway analysis for up and downregulated proteins are shown in Tabs 3 and 4, respectively (Microglia LPS Up KEGG; Microglia LPS down KEGG).

**Additional file 8: Table S6. LPS treated microglia phosphoproteomics, PPM1D^+/tr^ vs controls.** Phosphoproteomics was carried out on LPS treated microglia using the same samples described in Additional file 7: Table S5. Tab 1 (PPM1D+tr LPS vs Cont LPS) shows differentially phosphorylated phosphosites arranged by total score in descending order and the GO analysis. Tabs 2 and 3 (*PPM1D*^+tr^ LPS enriched Up KEGG; *PPM1D*^+tr^ LPS enriched Down KEGG) show the KEGG pathway analyses for phosphosites that increase or decreased, respectively.

