## Supplementary material for "Proteomics and phosphoproteomics profiling in glutamatergic neurons and microglia in an iPSC model of Jansen de Vries Syndrome": expanded methods

**Subjects**

The JdVS patient is a male who was born full term following an uncomplicated pregnancy. Oral motor weaknesses led to difficulty with feeding as an infant, requiring a referral with a feeding specialist. There was a delay in language development leading to speech/language therapy at 18 months of age. First word was spoken at 18 months. Receptive language was stronger than expressive language, although there were delays in the development of both. Hypotonia was noted and was managed by physical and occupational therapy. The patient walked at age 1 year 10 months. Medical history includes hypotonia, short stature, hyperopic astigmatism, and chronic otitis media with placement of pressure equalizing and adenoidectomy. He has a history of ADHD and severe anxiety, which are managed medically. A neuropsychological evaluation resulted in a diagnosis of intellectual disability. The Wechsler Intelligence Scale for Children - Fifth Edition (WISC-V) at age 9 showed a full-scale IQ in the 2nd percentile. A Differential Ability Scales – Second Edition (DAS-II) showed a General Conceptual Ability, Working Memory and Processing Speed in the 1^st^ percentile. The Neuropsychological Processing Concerns Checklist for SchoolAged Children & Youth – Third Edition (NPCC-3) showed good basic sensory function, motor function, and visual motor function. He has some difficulties with fine motor skills, such as buttoning and zippering, and has an aversion to loud noises. Math skills are poor. He is very friendly, sociable and polite. Whole exome sequencing at age 7 revealed a typical JdVS truncating mutation in exon 5 (c.1210C>T; p.Q404X). All *PPM1D* heterozygotes in this paper, whether patient-derived or developed using CRISPR-Cas9 editing (see below) will be referred to as PPM1D^+/tr^. Analysis of parental DNA showed that the mutation was de novo, as is the case for >90 of JdVS cases. The control for the patient-specific line is his typically developing older brother (by 3 years). The control used to develop isogenic iPSC lines with a truncation mutation in exon 5 using CRISPR was also a typically developing male used in another study (1).

**Development of iPSCs from peripheral blood CD34+ cells**

iPSC lines were generated from human peripheral blood CD34+ hematopoietic stem cells (HSC) with a CytoTune-iPS 2.0 Sendai Reprogramming Kit (Invitrogen) following the manufacturer’s protocol, as previously described (2). Frozen PBMCs were thawed 2 days before reprogramming (day -2) and cultured in STIF medium. On day 0, CD34+ cells were flow-sorted by FACSAria II (BD) and transduced with Sendai virus vectors containing Klf4–Oct3/4–Sox2, cMyc, and Klf4 in the presence of 4 µg/mL of Polybrene. Three days after transduction, the transduced cells were plated on a Matrigel-coated 24-well plate in StemSpan SFEM medium (STEMCELL Technologies). On day 7, half of the StemSpan SFEM medium was replaced, and on day 8, the culture medium was completely replaced. Thereafter, culture medium was changed every 2 days from days 2-7, then changed daily from day 8. The iPSC-like clones were picked and passaged by mechanical dissection from day 21 to day 28. FACS analysis of pluripotent markers (SSEA3, SSEA4, TRA-1-60, TRA-1-81), *in vitro* differentiation and immunohistochemical detection of 3 germ-layer markers (α-Fetoprotein, α- Smooth Muscle Actin and β-III Tubulin), RT-PCR assay for virus gene integration were performed on each iPSC clone to ensure that integration-free iPSCs with the capacity to differentiate into all three germ layers were generated. No clonal chromosomal alterations were detected.

**Differentiation of iPSCs into microglia**

To generate microglia, we used kits from STEMCELL^TM^ Technologies (STEMdiff^TM^ Hematopoietic Kit, catalog number 05310; STEMdiff^TM^ Microglia Differentiation Kit, catalog number 100-0019; STEMdiff^TM^ Microglia Maturation Kit, catalog number 100-0020) according to the manufacturer’s instructions, with minor modifications in the first step using the Hematopoietic Kit, during which iPSCs are conversted into HSCs. The iPSCs were grown in a 6-well plate until approximately 80% confluent, with spontaneously differentiating clusters of cells manually removed. For each well of cells, we prepared, in advance, a 12-well Matrigel plate, according to the manufacturer. When ready to start procedure, one ml of mTeSR Plus was added to each well of the Matrigel plate. The culture medium was aspirated from the iPSCs and were rinsed with PBS. One ml of detachment solution (0.5mM EDTA in PBS) was then added and the cells were incubated at 37oC for 3 minutes. Timing is crucial at this step. The detachment solution was quickly aspirated, and the wells were carefully rinsed with PBS, followed by the addition of 2mls of mTeSR Plus. At this point some cells might start to peel off, so the plates should be handled gently. Under a dissecting microscope, a StemPro EZPassage stem cell passaging tool (Invitrogen catalog #23181-010) was used to make horizontal and vertical cuts to produce 100um squares. The plates were then gently mixed to detach iPSCs. A 10ul pipette tip was used to collect individual squares from the plate (approximately 8ul, which contains 40-80 iPSC aggregates). Eight microliters containing aggregates was transferred to one well of the 12-well matrigel plate. After transfer, the cells were visualized under a dissecting microscope to ensure proper cell density and cell size. Once the iPSCs have been tranfered to the matrigel plates, the remainder of the HSC differentation protocol was followed according to the manufacturer, with one additional exception; at day 7 and day 10, the cells are fed by topping off with 0.5ml medium rather than removing entirely and replacing with fresh medium. On day 12, quality control was carried out by FACS analysis to show that populations expressed CD34, CD43, and CD 45 markers. Microglia Differentiation then continued according to the manufacturer’s instructions.

**Fluorescence-activated cell sorting (FACS)**

Single cell suspensions were used for flow cytometry staining. We followed a protocol for Staining Cell Surface Targets for Flow Cytometry from ThermoFisher. All antibodies were obtained from Stemcell Technologies, except for TMEM119, which is from Novus Biologicals. For hematopoietic stem cells (HSCs), we used CD45 FITC (Catalog number 60018FI.1) CD43 APC (Catalog number 60085AZ.1), and CD34 PE (Cat. 60013PE.1) antibodies. For microglia we used TMEM119 APC (Catalog FAB10313A) and CD11b PE (Catalog 60040PE.1) antibodies. Antibody concentrations were 5ul per 100ul for all Ab except TMEM119 for which 0.5ul per 100ul was used. Flow cytometry acquisition was obtained using a BD LSRII analyzer, and BD. FlowJo software was used for data analysis.

**CRISPR-Cas9 gene editing**

A heterozygous truncating variant in *PPM1D* exon 5 (*PPM1D*^+/tr^) was generated by CRISPR-Cas9 gene editing**,** using a protocol described by Ran et al {{5631 Ran,F.A. 2013}}. Briefly, a guide RNA (gRNA) sequence coding for a region in exon 5 adjacent to a PAM sequence 9 base pairs from the patient mutation the was chosen (see **Figure 1** in manuscript: 5’-ATAATAGTCAAGAAACCTGT). A 4 bp adapter sequence (5’-AAAC) was added to the 5’-end to facilitate cloning into the Bbs1 site of pSpCas9n(BB)-2A-Puro (Addgene catalog # 48139), which contains the Cas9 coding elements. A “C” nucleotide was added at the 3’ end (corresponding to a “G” nucleotide on the corresponding strand), which facilitates expression of the gRNA. The final gRNA sequence is 5’ AAACATAATAGTCAAGAAACCTGTC. This was annealed to it corresponding single stranded DNA (5’ CACCGACAGGTTTCTTGACTATTAT). The double-strand oligonucleotide was ligated into the linearized, Bbs1 digested plasmid. which were annealed prior to ligation into the linearized plasmid. iPSCs cells from a typically developing control were cultured and fed daily in mTeSR1 (Stem Cell technologies) on Matrigel (BD) coated plates at 37^o^C/5% CO_2_/85% in a humidified incubator. Cells were maintained in log phase growth and differentiated cells were manually removed before starting the experiment. iPSCs were exposed to 10uM ROCK Inhibitor for ~4 hours to improve cell survival during nucleofection. After 4 hours, growth medium was aspirated, and the cells were rinsed with DMEM/F12. iPS cells were dissociated into single cells with accutase and harvested by centrifugation. Nucleofection was performed using the Amaxa-4D Nucleofector Basic Protocol for Human Stem Cells (Lonza) according to the manufacturer’s instructions. Briefly, 8x10^5^ cells and 5ug of plasmid were nucleofected using the P3 Primary Cell 4D-Nucleofector X Kit L with program CA-137. Cells were resuspended in mTeSR1 + 10uM ROCK Inhibitor and placed in one well of a 6-well Matrigel-coated plate. The following day, cells were fed with fresh mTeSR1, and were subsequently fed with fresh medium daily. On days 4-14, cells were exposed to 0.5ug/ml puromycin for 6 hours. Puromycin-resistant colonies were picked and expanded in mTeSR1 without further puromycin treatment. DNA was analyzed by sequencing the region of interest. A PCR product flanking the engineered site was generated with the primers, 5’-TGTAGTGGCAGCTAAATCTGAG and 5’- CCAGGTGACGCTAACCAAAG, followed by Sanger sequencing. Truncating mutations were characterized using a cDNA translation tool to confirm a predicted frameshift mutation and the generation of a truncated protein https://web.expasy.org/translate/.

**Quantitative real time PCR (qPCR)**

qPCR was carried out on reverse transcribed PCR using the 2^-ΔΔCt^ method as we previously described (3-5).

**Western Blotting**

Proteins were prepared with PierceTM RIPA Buffer (Thermoscientific cat# 89900) according to the manufacturer’s protocol, with protease inhibitor cocktail mix (Sigma cat#P8340). Protein concentrations were verified using the BCA assay. Western Blotting was essentially carried out as previously described, with modifications. Briefly, 10-50 ugs of protein were denatured with the addition of Laemmli buffer and 2-mercaptoethanol, and boiled for 5 minutes. Samples were loaded onto a 12% precast polyacrylamide gel (BIO-RAD cat#456-1044). Gel electrophoresis was set at constant voltage (50V) for the first 30 minutes and 120V for the remainder of the run. The running buffer was in 1X TrisGlycine/SDS buffer. After separation by electrophoresis, proteins were transferred using the Trans-Blot^®^ TurboTM Transfer System according to the manufacture’s instructions. A seven-minute transfer was executed using the turbo program setting. After transfer, membranes were blocked in 5% milk with gentle agitation for one hour at room temperature. Membranes were then incubated overnight with gentle agitation at 4^o^C with primary antibodies for 24 hours. Cyclophilin B was used as a loading control. Primary antibodies used for WB included, Phospho-ATM Recombinant Rabbit Monoclonal Antibody mix (Cell Signaling catalog # 9607), PPM1D Polyclonal antibody, Rabbit/IgG, (Proteintech, catalog # 26532-1-AP), Cyclophilin B Polyclonal Antibody (Invitrogen, catalog # PA1-027A).

Following primary antibody incubation, membranes were washed three times with gentle agitation in 1X TBS/T buffer (20mM Tris Base, 0.136M NaCl, 0.1% Tween-20). Membranes were then incubated with a secondary antibody (1:5,000 dilution) plus anti-biotin (1:2,000 dilution) for 1 hour at room temperature with gentle agitation. Membranes were washed again, as above, and subsequently incubated with SuperSignal^TM^ West Dura Extended Duration Substrate (Thermo Scientific cat# 34075) for 5 minutes at room temperature with gentle agitation. Immediately thereafter, membranes were exposed to blue autoradiograph film for visualization. For CaMKII phosphorylation analysis, the relative expression of CaMKII was determined by taking the ratio of the autoradiographic signal of CaMKII and dividing it by the cyclophilin signal. The same was done for phosohorylated CaMKII using a specific Ab. Then, the relative signal for phosohorylated CaMKII was divided by the total CaMKII signal.

MAP2 (2a+2b) ; sigma cat# M1406, 1:500

VGLUT2 ; Millipore cat# MAB5504, 1:200

rabbit Anti-b-tubulin III ; GeneScript cat# A01203, 1:500

CAMK2 beta gamma delta (phospho T287), Abcam cat# ab182647; 1:500

CAMKII beta Abcam cat# ab34703; 1ug/ml

**Proteomics**

Proteomics analysis was performed as previously described (6,7). Briefly, cell pellets were lysed with 5% SDS in 50 mM triethyl ammonium bicarbonate, which results in a more efficient protein extraction and accession for digestion. Samples were treated with protease inhibitors, and all lysis steps were carried out on ice. Samples were reduced with 5 mM dithiotreitol for 30 min at 56 ^o^C and then carbamidomethylated with 20 mM iodoacetamide for 30 min at room temperature in the dark. Samples were then loaded onto the commercial cartridge S-trap (ProtiFi). The S-trap guarantees the removal of SDS prior to mass spectrometry analysis. Samples were digested with trypsin, and peptides were eluted from the S-trap. The peptide fraction was resuspended in 0.1% TFA and desalted using Oasis HLB C18 resin (Waters) loaded on 96-well filter plates (Orochem) mounted on vacuum manifold for 96-well plates. The peptides were washed with 0.1 % TFA and subsequently eluted with 60% Acetonitrile/0.1% TFA. Samples were subsequently lyophilized and stored at −80 °C for further analysis via mass spectrometry. Mass spectrometry was performed using nano liquid chromatography for high sensitivity and high performing state-of-the-art mass spectrometers (Orbitrap Fusion Lumos and Orbitrap Exploris 480, Thermo Scientific). Each sample was run with a three-hour gradient, which ensures exhaustive identification of the peptides separated by chromatography. Samples were acquired using data-dependent acquisition (DDA), which was used to generate the list of all peptides present in the samples. This hybrid acquisition method is currently the state-of-the-art for label-free quantification in mass spectrometry, as it ensures the lowest number of missing values and has a quantification accuracy comparable to targeted analyses (8). Peptides were identified by searching the DDA runs using Proteome discoverer v. 2.5 software. Data was transformed, normalized, and processed (7). Significance was assessed for p-values <0.05 and for fold changes >2 or <0.5. Each cluster was subjected to Gene Ontology enrichment analysis and the KEGG database**.** Differentially expressed proteins (DEPs) were rank ordered based on a “score” derived by multiplying the absolute PPM1D/control ratio by the p-value, the latter of which was calculated by a two-tailed heteroscedastic t-test comparing the relative abundances of the proteins between the two conditions. The p-value was then converted into -log2 to facilitate the plotting of the volcano plot. Enrichment of DEPs was determined using a web-based application called Gorilla, DAVID and EnrichR, which identify gene ontology (GO) terms based on a ranked list of genes or proteins based on their degree of differential expression (11).

**Phosphoproteomics**

Cell pellets were homogenized as described above. Following S-trap elution, phosphopeptide enrichment using titanium dioxide (TiO2) chromatographic resin was performed as previously described (12,13). The peptide samples were subsequently lyophilized and stored at −80 °C for further analysis via mass spectrometry. The lyophilized phosphorylated peptide samples were reconstituted in 0.1% trifluoroacetic acid (TFA) and analyzed using nano liquid chromatography tandem mass spectrometry (LC-MS/MS) as described above. Both the proteome and the phosphoproteome were analyzed for each sample; the proteome was used to normalize the phosphorylation changes by the protein level changes, to ensure that the observed regulations are due to the actual level of the phosphorylation. Samples were acquired using data-dependent acquisition (DDA) and MS/MS data were used in the subsequent automated protein database search to generate the list of all phosphopeptides present in the samples. We used a two-step normalization: first, phosphopeptides were normalized by the center of the distribution of their abundance to correct for imprecisions in sample injection amount; then, we normalized them by protein abundance changes, to correct for protein regulations that might be misinterpreted as regulation of phosphorylations. Normalized values were compared by using parametric statistic after passing the Shapiro-Wilk test, which evaluates whether data are normally distributed. Due to multiple conditions, we used paired analyses using the t-test. Significance was assessed for p-values <0.05 and for fold changes >2 or <0.5.

To simplify data interpretation, the significantly regulated phosphorylations were displayed using Microsoft Excel.

stylefix

Works Cited
