## Supplementary figures and images for "Proteomics and phosphoproteomics profiling in glutamatergic neurons and microglia in an iPSC model of Jansen de Vries Syndrome"

### microglia in suspension

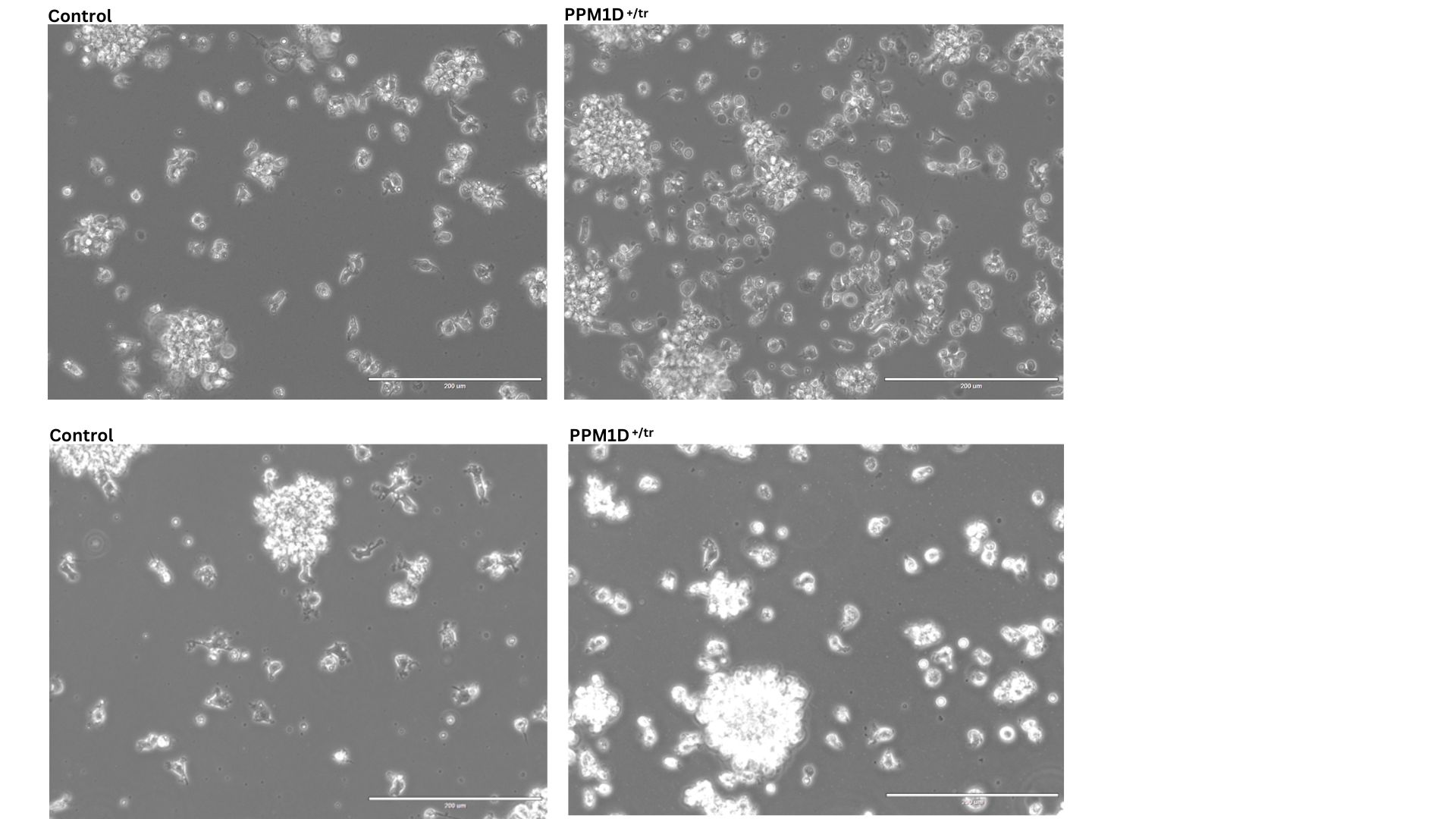
